# Multiscale cosimulation design template for neuroscience applications

**DOI:** 10.1101/2022.07.13.499940

**Authors:** Lionel Kusch, Sandra Diaz, Wouter Klijn, Kim Sontheimer, Christophe Bernard, Abigail Morrison, Viktor Jirsa

## Abstract

Integration of information across heterogeneous sources creates added scientific value. It is, however, a challenge to progress, often a barrier, to interoperate data, tools and models across spatial and temporal scales. Here we present a design template for coupling simulators operating at different scales and enabling co-simulation. We illustrate its functioning along a neuroscience example, in which individual regions of interest are simulated on the cellular level to address mechanistic questions, while the remaining network is efficiently simulated on the population level. A workflow is illustrated for the use case of The Virtual Brain and NEST, in which the cellular-level hippocampus of the mouse is embedded into a full brain network involving micro and macro electrode recordings. This new tool allows integrating knowledge across scales in the same simulation framework and validate them against multiscale experiments, thereby largely widening the explanatory power of computational models.

## Introduction

The brain is a complex system that includes billions of cells that interact with each other in a nonlinear manner. As a result, even if we were able to measure what all cells are doing simultaneously, we would not gain a deep understanding on how the brain works. Theoretical models can account for nonlinearities and emergent properties. Numerous models have been developed to study molecule interactions inside cells, cell physiology, activity of cell populations, up to body behaviour [1,2]. It is currently impossible to model the brain with all its cellular and molecular constituents due to limitations in resolution, computational resources, or available data from measurements. As a result, even if a given physio/pathological process can be modeled at the macroscopic scale, the lack of microscopic resolution at the molecular scale prevents obtaining mechanistic insight [3]. It is therefore important to bridge different scales. In material science, the study of composite materials requires the description of molecular interactions of individual composites, and a global description for the analysis of the subsequent deformation of the composite plate [4]. In biology, to understand the effect of drugs on tumour growth, it is necessary to model the tissue of cells around the tumour, the tumour cells, and the subcellular transduction signalling pathways [5,6]. In neuroscience, synaptic plasticity uses mechanisms of spike timing on the millisecond scale but leads to the formation of long-term memory evolving on the scale of minutes, days and weeks [7]. The goal of this study is to provide a methodology to address scientific and technical problems of multiscale co-simulation applied to the brain.

The main difficulty of multi-scale modelling is coupling the different scales. This coupling cannot be generic because it depends on the model and the properties of the data. For example, in the case of tumours, the tissue around the tumours is represented by a continuum model (first scale), which interacts with discrete tumour cells (second scale); while continuous signalling pathways are modeled in cells (third scale). At present, it is not possible to create a common coupling function between these three scales.

Another difficulty lies in the fact that different simulator engines have been developed to study a system at a given scale. In the case of tumours, a common approach is to use COMSOL Multiphysics [8] for the tissue simulation and Matlab [9] for the cell and subcellular scales. Since most simulators do not include external interactions, it is difficult to link them within a common framework.

Furthermore, it is not possible to extend existing solutions developed in physics [10,11] or in biology [12,13], due to the specificity of the simulators, models and emergent properties at lower scales. In particular, neuroscience has multiple models for describing neuronal activities, and each scales has multiple simulators for running these models (e.g. neuronal level: Neuron [14], Arbor [15] Genesis [16]; network level: NEST [17], Brian [18]; system level: The Virtual Brain (TVB) [19], Neurolib [20]). Some multiscale models using co-simulation make use of the existing solutions [21–24] but there is no co- simulator for point neuron network models and whole brain network models. Such co-simulator would enable a better understanding of why individual neurons fire action potentials given the state of whole brain activity. Such a framework is important to interpret experimental data combining microscopic Local Field Potential (LFP) and neuronal firing recordings, and macroscopic electro-COrticoGraphy (ECOG) in mice [25].

Here, we present a methodology to address the problems of multiscale co-simulation. The method is based on a generic design template which dictates a separation of science and technical attributes, allowing these be addressed in isolation where possible. This separation is based on transformer modules, which are used for the synchronization and connecting simulators and they also include the function for transforming data between scale. This method is applied to construct a multiscale model from experimental data obtained in the mouse brain with ECOG cortical signals and LFP signals in the CA1 region of the hippocampus. This model is running on a co-simulator prototype using the simulators TVB and NEST. Three sets of parameters of this model, resulting in different network dynamics, are chosen in order to demonstrate the feasibility and the limits of this modelling approach. The following sections describe the technical details and the optimizations required for coupling TVB and NEST as a way to illustrate one prototype implementation of the template and to identify its technical limitations.

## Results

The multiscale co-simulation design template formalizes the interactions between parallel simulations at different scales. The transformation of the data among scales is performed during their transfer among simulators. This template is composed of 5 modules (figure 1a): one launcher, two simulators, two transfer modules. Each transfer module contains 3 components : one interface for receiving data, one interface for sending data and a transformation process. The launcher starts and handles the coordination of simulation parameters. The simulators perform the scale specific simulations. The transfer modules transfer the data from one simulator to another. During the transfer, the transformation process transforms the incoming data for the simulator on the receiver side.

**Fig. 1.**
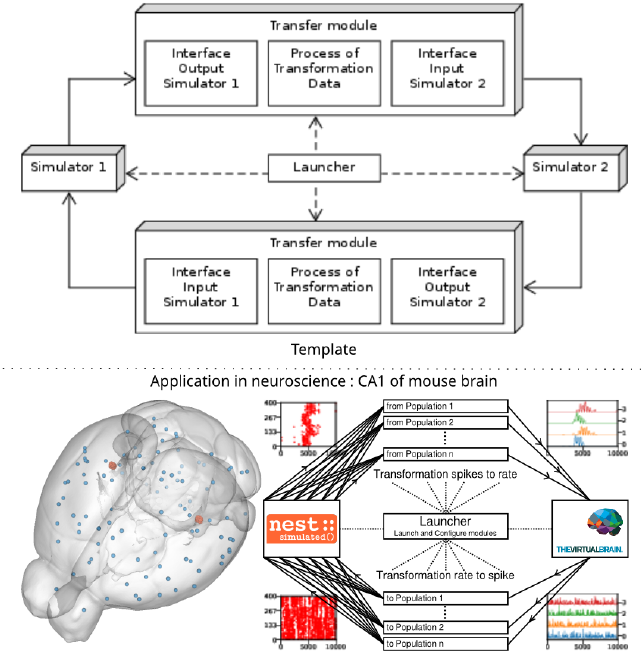
Multiscale co-simulation design template and example of application in neuroscience Top. Multiscale co-simulation design template between 2 simulators using transfer modules for transformation and transfer of data between scale. **Bottom**. Application of the co-simulation template for a neuroscience use case focusing on the CA1 region of mouse brain. The **left panel** shows a rendering of the mouse brain from Allen Institute[26]. Blue spheres mark the centres of mouse brain regions and the red spheres a subset of neurons of the CA1. The **right panel** illustrates the co-simulation data flow between TVB and NEST showing the different functional modules. The plots in the four corners illustrate the type of data exchanged in respective information channels. The transfer modules exchange mean firing rate data with TVB (module on the right) and exchange spike times with NEST (module on the left). Each population has a specific module enabling transfer of data between populations in different scales.

In this study, the multiscale co-simulation template is applied to a virtual experiment workflow between the in-silico mouse whole-brain dynamics and the in-silico micro-scale network dynamics of the hippocampus CA1 region. The recording of the virtual CA1 and virtual mouse brain as similar than experiments[25] (see figure 1b). The Virtual Brain (TVB)[19], an open source platform, has been used to simulate the mouse whole-brain network activity, while NEST[27], another open-source platform, has been employed for the simulation of the CA1 neuronal network dynamics. This specific application is used to illustrate the technical limitations of this novel design template and to demonstrate the potential for a wider range of applications.

### Virtual experiment of hippocampal CA1 embedded in a full mouse brain

The virtual experiment of the mouse brain is composed of an brain network model, regional neuronal network models and electrophysiological sensors models.

The whole-brain animal model is a network comprised of nodes and edges, where each node contains a neural-mass model to simulate the activity of each region and where edges represent the anatomical connections among the regions. The anatomical connections are defined by track lengths and an adjacency matrix representing the coupling strengths of connections between the regions of the network, the “connectome”, which are extracted from tracer data from the Allen Institute[29] (figure 2f and 2g). The dynamic activity of each brain region is obtained with the neural mass model described by Di Volo et al. paper [30] (see Online Methods). The neuroinformatics platform The Virtual Brain (TVB)[19] is able to perform the animal whole-brain simulation by considering both the chosen neural-mass and the specific “connectome”. The dynamics of the two main brain regions of interest, the left and right hippocampus CA1 (figure 2), are modelled as a separate neural network composed of point neurons connected with static synapses. Each network is composed of one inhibitory and one excitatory homogeneous population of adaptive exponential integrate and fire neurons [31] (see Online Methods). In each microcircuit, the populations of point neurons are taken to be homogeneous, that is, neurons of the same population have the same parameter values. The neuroinformatics platform NEST[27] is able to perform the regional neuronal network simulation using the aforementioned description of the microcircuit of point neurons.

**Fig. 2.**
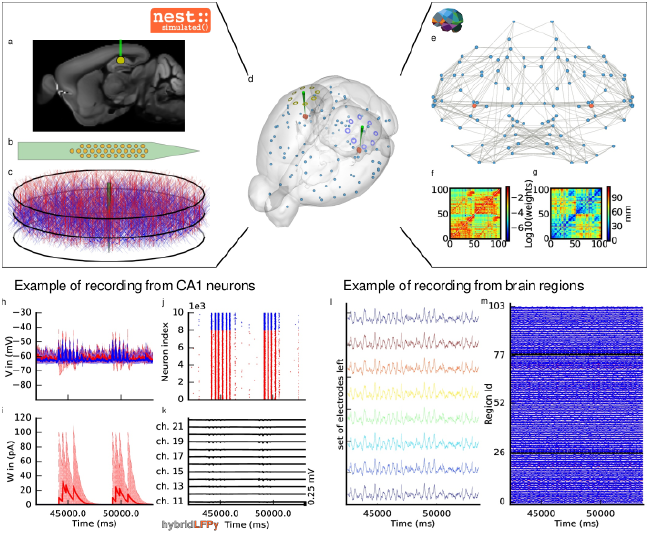
The virtual mouse brain experiment. **a** Cross section of the mouse brain with the position of the left implanted electrode. **b** Position of the site layout of the polytrode (Neuronexus 32 models from MEAutility library). **c** The position of the probe inside the neural network. The red neurons are pyramidal neurons[28] and the blue neurons are basket cells[28]. **d** Mouse brain of Allen Institute[26] with the position of the 2 polytrode electrodes and 16 ECOG electrodes. The ECOG electrodes measure the neural field from the surface of the electrode in blue for left hemisphere and yellow for the right hemisphere. Blue spheres mark the centres of mouse brain regions and the red spheres a subset of neurons of the CA1. **e** Representation of the connectome of the mouse brain[29]. The blue dots are brain regions and the red ones are CA1 regions, whose neurons are simulated with NEST. The strongest anatomical connections are highlighted by the grey links. **f** The weights of the anatomical links in F are shown as an adjacency matrix. **g** The tract lengths associated with F are shown as an adjacency matrix. The anatomical connections are extracted from tracer data of the Allen Institute[29]. **h** example of voltage recorded from 10 excitatory and 10 inhibitory neurons. **i** Example of adaptation currents recorded from 10 inhibitory neurons and 10 excitatory neurons. **j** Example of spike trains recording from the left CA1. **k** Example of local field potential recorded from the poly-electrode which is generated from the spike trains and the neuron mythologies. **l** Example of recording from the ECOG electrodes of the left hemisphere. **m** Example of mean firing rate of excitatory and inhibitory population each region of mouse brain

In order to compare the simulations with empirical data, the virtual experiment contains two models of electrophysiology sensors for probing neural activity. The electrophysiological sensor models are two surface grids composed of 8-channel electrocorticography arrays and two penetrating multi-electrode arrays composed of 32 recording sites each. Their positions are illustrated by the figure 2a. The figure 2e shows the position of the polytrodes in the mouse brain, while the figure 2b and 2d depicts the position of the left probes in a cross-section of the left hemisphere and the position of the point of the polytrodes in the population of neurons, respectively. Figure 2c displays the polytrodes with the 32 recording sites. The simulated signal from the ECOG sensor is computing using the model of a point dipole in a homogeneous space as described by Sanz-Leon et al. 2015[32] (see Online Methods) and the hybridLFPy[33] software is used for computing the signal from the recording site of the implanted probes (see Online Methods). The latter software uses morphology and spatial position of neurons to generate the underlying local field potential (LFP) for given spike trains of point neurons. The morphology of the excitatory neurons is taken to be that of the morphology of pyramidal cell from the model in Shuman et al. paper[28] (see Online Methods). From the same model, the morphology of a basket cell is taken for the morphology of the inhibitory neurons.

### Output signal from the virtual experiment

This section describes the co-simulation results at different scales by describing the possible recording physiological signals of the virtual CA1 in full-mouse brain. The Discussion section will provide interpretation of these results for describing the advantages and the limitations of the multiscale co-simulation template. As described in figure 2, the output modalities of one virtual experiment are comparable to the outputs of a real experiment which are the local field potential measure at each thirty-two sites of each polytrode electrodes (figure 2j) and from the sixteen electrocorticography channels of each hemisphere (figure 2k). Moreover the simulation gives access directly to the voltage membranes of the CA1 neurons (figure 2h), adaptive current of the CA1 neurons (figure 2g), spike times (figure 2i) and the mean firing rate of the different regions of the mouse brain (figure 2m). For an illustration of the possibility, a set of three different parameters of CA1 is chosen. Each parameter represents one of three dynamic regimes of the CA1. This results are separated between micro (figure 3) and macro (figure 4) scale but they are the output of the simulation workflow between TVB and NEST. In particular, figure 3 reports the mean voltage membrane, mean adaptive current, instantaneous firing rate and the signal of 12 central sites from the 32 electrode sites of the specific CA1 network. Figure 4 displays the results on the whole brain level: the mean firing rate of each brain region, the signal of the 16 electrocorticography channels and the transferring mean firing rate from the spiking neural network.

**Fig. 3.**
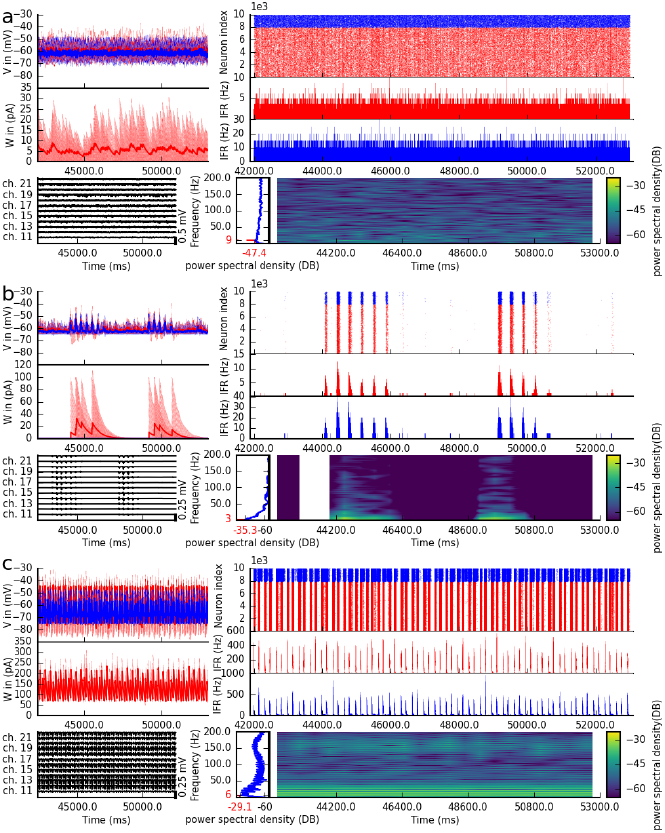
Spiking neural network in 3 different states of the left CA1. The parameterization of the spiking neural network of CA1 is chosen such that the dynamics are in a asynchronous state (**a**), irregular synchronization state (**b**) and regular bursting (**c**). **top-left** Voltage membrane of 20 adaptive exponential leaky and integrator neurons and the mean of them in thick line. The red (blue) lines are excitatory (inhibitory) neurons. **middle-left** The adaptation currents of 10 neurons are shown and the mean of them in thick line. **bottom-left** Local field potential from the 12 sites of in the middle line of the polytrode. The local field potential is computed from the spike trains of all neurons by the software HybridLFPY[34]. **top-right** Spike trains of 10000 neurons for 11s. **middle-right** instantaneous firing rate of the excitatory (inhibitory) population above in red (blue). **bottom-right** Spectrogram and power spectrum example of the instantaneous firing rate for 10s.

The figures 3 and 4 are separated into three different panels, which correspond to the three sets of parameters representative of the different types of dynamics exhibited by spiking neural networks (see online method for the choice of these parameters). Panel **a** represent an asynchronous (A) state, which is characterized by a constant (flat line) the mean firing rate (see figure 3a top-right). Panel **b** represents an irregular synchronous (IS) state which reflects large irregular variation of the mean firing rate (see figure 3b top- right). Panel **c** represents regular bursting (RB) reflecting regular oscillations (see figure 3c top-right) and a second dominant high frequency (see figure 3c bottom-right).

#### Results at microscale

The top left of the panel a, b and c of figure 3 show the membrane voltages for ten excitatory neurons (thin red curves) and ten inhibitory neurons (thin blue curves) and mean membrane voltage of these neurons (thick curves). The middle left of the panel a, b and c of the figure 3 represent the adaptive currents from the same ensemble of neuron (thin curves) and mean adaptive current of these neurons (thick curves). The third biological observable from the simulation is the Local Field Potential which differs among panels (see bottom left of panel a, b and c of figure 3). The top right of panel a, b and c of figure 3 display spike raster plots of the excitatory population, in red, and the inhibitory population, in blue, of the left CA1. The spiking activity is homogeneously distributed between neurons and time frames for the A state, while the other two states show co-activation of neurons with different periods. The associated instantaneous firing rate is shown in middle right of panel a, b and c of figure 3. The spectral analysis of the instantaneous firing rate displays a peak around 3 Hz for the IS state (bottom left of panel b of figure 3), no peaks for the A state (bottom left of panel a of figure 3). and two peaks (around 6 Hz and 160Hz) for the RB state (bottom left of panel c of figure 3). For the RS state, the frequency of the first peak, 6Hz, is also present in the mean of the adaptive currents while the second peak is associated with the burst time, as shown with further detailed in the Supplementary Figure 1.

#### Results at macroscale

The top left of the panel a, b and c of figure 4 display the instantaneous firing rate (light red) of the spiking neural network with the associated transferred mean firing rate of the left region of CA1 (thick red line). The ECOG signals are affected by the different states of the neural network, as shown in the bottom left of panel a, b and c of figure 4. The mean firing rate of excitatory (blue) and inhibitory (red) population of each brain region are plotted in the graph on the right part of panel a, b and c of figure 4).

**Fig. 4.**
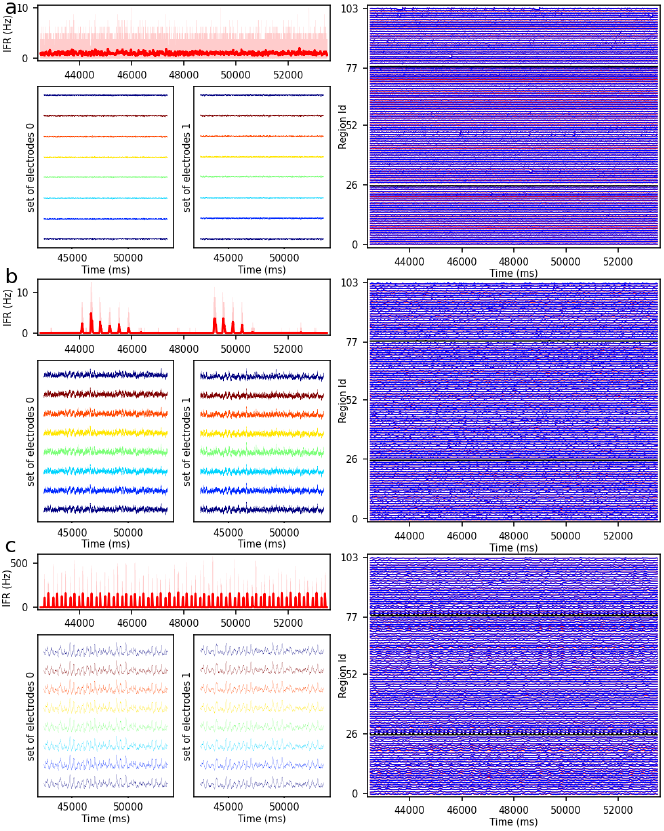
Three different states of CA1 in mouse brain. The parameterization of the CA1 spiking neural network is defined to obtain asynchronous state (**a**), irregular synchronization state (**b**) and regular bursting (**c**). **top-left** Instantaneous firing rate of spiking neural networks in light red for 11 second time span. The thick line shows the sliding window mean firing rate. **bottom-left** (**bottom-right**) Signal from ECOG sensors, the figure is for recording of the 8 electrodes on the top of the left (right) hemisphere. **right part** Full region overview of the mean firing rates of excitatory, in red, and inhibitory, in blue, population from the model of Mean Adaptive Exponential. The 2 black curves are the mean firing rate of the 2 population of excitatory neurons simulated with NEST[27].

### Workflow between NEST and TVB

The previous multiscale example uses the workflow between TVB and NEST for the co-simulation. This workflow, as an implementation of the design template, is composed of five modules : two simulators (TVB and NEST), one launcher and two transfer modules. All these modules built with the capability to be repurposed or replaced allowing for adjustments of components of transfer module or communication protocols (see Discussion). For demonstrating the possibility of reusability of the components, two additional proofs of concept were implemented. The first one replaces NEST by NEURON and the second one replaces TVB by Neurolib (see Supplementary Figure **??**). Moreover, without extra development, we get a proof of concept of co-simulation between NEURON and Neurolib.

The simulators perform the actual integration of the dynamics in time and require two properties to be integrated within one optimized and coherent workflow. The first property is the time delay equation management which is essential for reducing data transfer overhead. The second property is the presence of a high bandwidth Input/Output (I/O) interface that facilitates the efficient exchange of data and parallel execution of the simulators. Since TVB and NEST did not have generic high bandwidth I/O interfaces by default, these had to be implemented for each simulator. Details of how these I/O interfaces were created are reported in the Supplementary Note 1. Briefly, the NEST interface uses the device nodes with a specific back-end, while TVB uses proxy nodes which are proxy nodes used for the interface with the external software.

The launcher prepares the environment for the simulation and initiates all the other modules, as shown in the figure 5a (see details in the Supplementary Figure **??**). The preparation consists of the creation of folders for the different modules as well as for the logger files and the common file with all the parameters of the co-simulation. The creation of the parameters file provides the functionality to enforce consistent constraints on the parameters which are to be shared between the modules, such as ensuring the use of the same integration step in both simulators, which is needed for correct synchronization between modules.

**Fig. 5.**
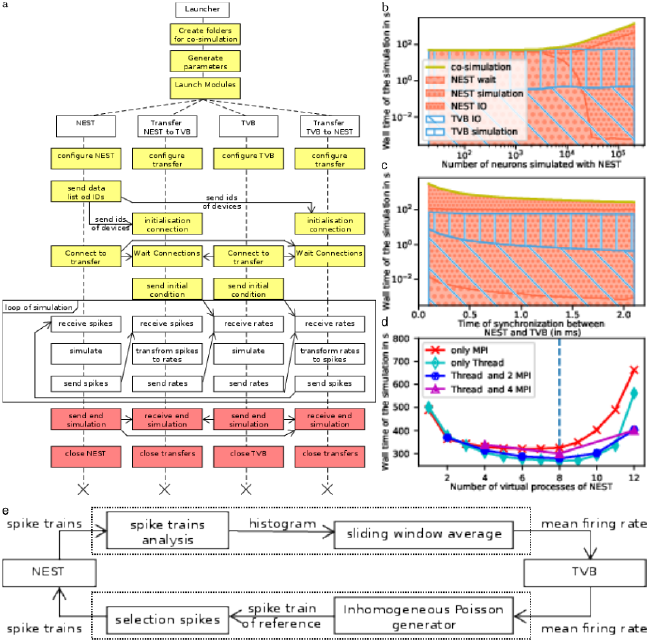
Architecture and performance of the co-simulation. **a** The interaction among the modules and data exchanges during co-simulation execution. The boxes in yellow mark start-up:initialization and configuration, the boxes in red the termination of the simulation and the boxes in white for the simulation phase. **b**,**c**,**d** Performance of the workflow is obtained for 1 second of simulated biological time (see Online Method for more details). The reference implementation use 1 MPI process, 6 virtual processes/threads, a synchronization time step of 2.0 ms, and simulates 20000 neurons. **b** The wall clock time of the simulators as a function of the number of neurons. The total time of the co-simulation is represented in yellow. The “wait”, “simulation” and “IO” times of NEST are represented in red surface with respectively hatches with big circles, small circles and points. The “simulation” and “IO” times of TVB[19] are represented in the blue surface with respectively hatches horizontal lines and oblique lines. **c** Simulation time depending on the synchronized time between simulator. The colour code is the same as the panel B. **d** Wall clock time depending on the number of virtual process used by NEST. The green, blue, purple, red curves are associated with different parallelization strategy of NEST, respectively, only multithreading, 2 MPI processes with threads, 4 MPI processes with thread and only MPI processes. The vertical blue line represents the number of cores of the computer. **e** The ‘transform between spikes to rate’ and ‘transform between rates to spikes’ blocks are displayed with the different steps for transformation of data between TVB and NEST.

The transfer modules are connecting simulators by transferring data between scales and by adapting the delay of communication throughout the simulation. Each module is comprised of three components : two interfaces and one transformer (see Figure 1a and Supplementary Figure **??**). These components are implemented in different files for reusability and modularity and are tested independently to ensure robustness (see Supplementary figure **??**). The interfaces are specific to each simulator while the transformation can be extended, modified or reused since the transformation function is implemented as an independent process (see Supplementary Note 2).

The different components use a simple Application Programming Interface (API) for the exchange of data between them. The API is implemented with two different technologies depending on the nature of the parallelization of the components (multiprocessing or multithreading). In the case of multiprocessing, each component runs in an individual process and a Message Passing Interface (MPI) is used for the transfer of data. In the case of multi-threading, each component runs in an individual thread in a shared process and the data is transferred using shared memory. The multithreading uses less computational resources (see Performance section) though it is not practical, specially on super computer, due to the global interpreter lock of python (for more details see Online Methods).

The transformation function provides mean field firing rate values by using a sliding window, shown in figure 5e. The panel also illustrates the inverse transformation from the mean firing rates to spike trains using a multiple interaction process[34].

The modular workflow execution is composed of three main blocks: start-up, simulation-loop and termination (see Figure 5a and details in the Supplementary Figure **??**).

The start-up procedure allocates a logger for each components facilitating easy debugging of the co-simulation. Subsequently, the modules and their communication channels are configured according to parameters file. At this stage a number of initialisation files are generated with simulation parameters only available after instantiation of the model (e.g. id of NEST devices and MPI port description).

Once the simulation is launched, the simulator time clocks are synchronized using asynchronous message passing: At each synchronization step, simulators receive input data after which the next step is simulated. The transfer modules can buffer data for one synchronization step until the the receiving simulator is available for receiving. Each simulator requires an initial condition (NEST : initial voltage membrane and adaptation current and TVB: state of the node during the previous seconds) and an initial message. For TVB, this starting message is sent by the transformer processes while, for NEST, it is produced by transforming the initial condition of TVB.

Ultimately, the termination occurs at the end of the simulation by the simulators themselves (see Online Methods for details).

### Performance

The evaluation of the performance is against a fictitious workflow with optimal performance, a co-simulation with instantaneous communications between simulators. The reason for this evaluation is due to the novelty of the multiscale co-simulation template stems from the presence of the transfer module that ensure coherence exchange between micro-scale and macro-scale. As all the modules are designed to run in parallel, the co-simulation time for each module is identical and equal to the total running time. The focus is only on the simulator timers because the time of the transformer components is dominated by waiting time of data (see Supplementary Figure **??**). The total running time of the simulators is divided in 5 parts. The “initialisation” time is the time of the configuration of the simulators and the creations of connections. The “ending” time is the time for simulators to close the connections, stop the simulator engine and terminate the processes. The “simulation” time is the total duration of the internal computation of simulator engines. The “wait” time is the total duration of waiting time for access to the data to transfer by the simulator interface of the transformer module. The “IO” time is the total duration of function for exchanging data between simulator and the transfer modules minus the “wait” time.

A perfect co-simulator has the time of the slowest simulator X, thus “wait” and “IO” times equal to zero. From the figure 5b and the Supplementary figure **??**, the actual implementation is closed to ideal when the number of neurons simulated by NEST is lower than 1000. In this case, TVB is the slower simulator and NEST spends most of the time waiting for data from TVB. When the number of simulated neurons is between 1000 and 20000 neurons, “simulation” time of TVB is approximately the same as the sum of “simulation” time and “IO” time of NEST. In this condition, each simulator is waiting for the transformation of the data among scales.

When the number of simulated neurons is higher than 20000, NEST is the slowest simulator. In this case, the co-simulation time is determined by the “simulation” time and the “IO” of NEST. The “wait” time is set to zeros and the “IO” time is higher than the “simulation” time (see Supplementary Figure **??** and Supplementary Figure **??**). The two principal causes are that the communication between modules is slower than inside the modules and the increase in size of the neural spike data with the increasing number of neurons (each neuron in NEST receive an individual spike train). With a closer look at the performance, we can see that the communication spends the majority of time on sending individual spike trains to NEST (see Supplementary Figures **??**). However, the size of data is related to the model chosen and can be reduced.

As displayed by the figure 5c and 5d, some optimizations can be implemented to reduce the problem of overhead time of communication. Figure 5c and Supplementary figure **??** represents the time delay between brain region when delayed data is aggregate to reduce the “IO” time and, hence, the co-simulation time. In this case, the simulators are not synchronized at each time step but at n time steps (limited by the model of connection). This aggregation can reduce co-simulation time by a factor 6 (see Supplementary Figure **??** and Supplementary Figure **??**). Figure 5d and Supplementary figure **??** represents a reduction of co-simulation time per reduction of the “simulation” time of one simulator. The increase of the resources used by NEST does not modify the “IO” time until the resource is available. Since the tests are running on one computer, the increase of resources for NEST increases the “simulation” time of TVB and reduces the “simulation” time of NEST. However, by deploying the workflow on high-performance computing facilities, the latter result does not replicate and the simulation time gives similar a result with an increase in “IO” and “simulation” time because the communication between nodes is slower (see Supplementary Figures **??**).

## Discussion

The paper presented co-simulation technology linking two simulators operating on two different scales together with minimal requirements and modifications. This workflow is based on cyclic coupling topology of modules[35] with generic coupling function for each scale. Nevertheless, the simulator of specific scale are interchangeable due to the modularity of the transfer module and the design of the template (for more characterization of the workflow, see the Supplementary Note 3). The interfaces of the simulators and other modules can be reused in other studies involving co-simulations. In comparison, MUSIC[24] requires extensive interface integration for communication with an orchestration and the standard like High Level Architecture[36] and Functional Mock-up Interface[37] does not accommodate the introduction of transformation modules. One important point is the integration of previously developed tools at different scales in the same workflow, which is essential for the validation of simulations against experimentation and the robustness of these future results.

Our current design template separates the theoretical problems of coupling models from different scales and the technical problems of coupling simulators. In addition, the template gives solution to other technical aspects such as the synchronization of simulators. Syntactic, semantic, and conceptual issues remain to be solved[44]. The design template allows communities working on different scales to work together for further evolutions of co-simulation models. The implementation of the template for a particular problems is a starting point to capture the interest of both communities by creating examples and to discuss the scientific part without being bogged down in technical details. However, the template doesn’t provide technical guidelines for the robustness, the management and the maintenance of the co-simulation. These challenges are the focus of the closely related staged deployment and support software for multi-scale simulations developed in EBRAINS (https://juser.fz-juelich.de/record/850819). Nevertheless, common mistakes in multiscale modeling are of conceptual nature, this paper contains two mistakes. The first mistake is not to apply the model following its definition. By definition, the neural mass model used in this paper cannot capture the fast scale, in particular, the fast regimes of regular burst state[38]. The second mistake is the non-respect of the assumption of the model. The neural mass model considers that the excitatory input firing rate of neurons is an adiabatic process. This hypothesis is broken in the irregular synchronous state due to the presence of rapid transition between low and high firing rates. Responsible use of multiscale models thus requires an understanding of models when operating the simulation engines. Numerical errors constitute another issue. As these errors cannot be estimated analytically, the solution is to perform a sensitivity analysis or uncertainty quantification to determine whether or not the simulation result is reliable[39][40].

For the validation of the co-simulation, it is important to generate data that can be related to real-world observations, as is the case here with the model of the two types of electrodes. An second important properties is the repeatability and reproducibility of the simulations. Repeatability is ensured by managing all the random generators in each simulator and using a single parameter file for the co-simulation setup. For reproducibility, due to the complexity of the network, a table is proposed where the configuration of each simulator is reported with their version and also the description of the transformation modules (see Supplementary Table 1). An important property of this design template is the independence of the modules and components. This independence allows unit tests to be performed for each of them, which is important for maintenance, debugging and robustness of the simulation. The actual workflow TVB-NEST provide a minimal approach for these three properties, which are essential for the long term development of the co-simulation framework. Our design model pushes developers to create a minimal interface of a simulator for its interaction with another one, because most of the time, simulators are meant to be run in isolation[41]. This interface must be reusable, to increase the number of users, which is important for its maintainability. This interface can be retrofitted to standards such as High Level Architecture[36], Functional Mock-up Interface[37], CAPE-OPEN Interface[42] or a future standard.

The workflow between TVB and NEST provides a new approach to address multi-scale problems. This workflow includes a direct link to experimentation, such as Opto-E-Dura[25] and in future work will be extended with functionality to directly analyse data recorded and compare them with experiment data. This workflow allows simulation of microcircuits while considering the interconnection with the rest of the brain. Furthermore, one advantage of this method in comparison with top-down or bottom-up approaches, it is the reduction of ambiguity when we change scale due to the explicit definition of transformation functions[1,3].

In summary, we have presented a new template for coupling simulators with a transformation module. This template provides the first step for the development of platforms using transitional scaling models and structure the future syntactic, semantic and conceptual issues induced by multi-scale problems. One optimization performed for this specific workflow is based on the communication delay between scales. It is not generalized for all cases but it is recommended for models with transmission line element method[43] or waveform relaxation method[44].

## Acknowledgments

This research has received funding from the European Union’s Horizon 2020 Framework Programme for Research and Innovation under the Specific Grant Agreement No. 945539 (Human Brain Project SGA3) and Specific Grant Agreement No. 785907 (Human Brain Project SGA2). The authors would like to thank Mario Lavanga for helpful feedback, Ingles Chavez Rolando for technical support and numerous colleagues for comments texts and figures. Also, the authors gratefully acknowledge the support team of the supercomputer DEEP-EST and JURECA at Forschungszentrum Jülich and Piz Daint at CSCS - Swiss National Supercomputing Centre.

## Code availability

The co-simulation between TVB and NEST is freely available under v2 Apache license at https://github.com/multiscale-cosim/TVB-NEST on the branch Paper TVB-NEST and Paper TVB-NEST with timer.

A docker container which contains the project is freely download at on Ebrains (https://docker-registry.ebrains.eu/harbor/projects/53/repositories). The simulated data of the three set of parameters, the code and the singularity images are available in a zenedo repository (http://dx.doi.org/10.5281/zenodo.7259022).

## Online Method

The details of the simulation and parametrization of the models can be found in the Supplementary Table 1. The format of this table is drawn from the proposition of Nordlie et al. 2009[1]. The proposition of Nordlie et al. is for spiking neural networks. This new format includes the description of brain network modelling, the description of the coupling between scale and the description of the measurements of the simulation. This format contains more details than the proposition of Nordlie et al. because it contains all the parameters for the co-simulations.

The following text provides an overview of the models, communication between modules, details of the performance tests and implementation details.

### Models

#### CA1 model

The spiking neural network of CA1 is comprised of two regions (left and right), which contains two populations, 8000 excitatory neurons and 2000 inhibitory neurons. This network is simulated by NEST[2], a neuro-informatics platform for spiking neural networks. The adaptive exponential integrate and fire neurons[4] are connected by exponential conductance-based synapses with a connection probability of 5% in side the region. The excitatory population establishes connections among regions in fixed amounts defined by the mouse connectivity atlas and the fixed number of synapses received by each neuron from other regions. The delay of transmission between regions is defined as the ratio of distance between the regions and the speed of transmission. The calculation of these ratios are part of the configuration of The Virtual Brain (TVB)[19] because the data required by TVB is the track lengths between regions and the speed of the transmission. Within a region, the synaptic transmission delay is instantaneous. In addition, the neurons can receive external noise input modeled as an independent Poisson process in addition to the external stimuli received from other regions through the transfer of mean firing rates as transformed spike trains.

#### Mouse Brain model

The mouse brain model is simulated using The Virtual Brain[19] which is a neuro-informatics platform for connectome-based whole-brain network modeling. The “connectome” used here is extracted from Allen Mouse Brain Connectivity Atlas[6] in 2017. The large-scale brain network is comprised of linearly coupled neural mass models. Specifically, the model representing each region is a second order Mean Ad Ex[7] with adaptation, which represents the mean firing rate for an ensemble of one excitatory and one inhibitory population neurons.

#### Electrophysiological monitoring model

The electrophysiological monitoring variable are computed using 2 models, representing the cortical and implanted sensors. The electrocorticography model is a forward solution of a dipole at the region level. Each of the brain region is reduced to its centre and the dipole is considered in a homogeneous space[8]. The implanted sensors signal are computed from point-neuron activities using a hybrid scheme for modelling local field potentials (LFP). Specifically, each potential is simulated using hybridLFPy[9] which incorporates the recorded spike from the network and the morphology of the pyramidal and basket cells.

#### Choice of three set of parameters

The parameters for Irregular Synchronous state are chosen based on Di Volo et al. paper[7]. The coupling between region and the noise was define after empirical exploration in order to get a fluctuation of the firing rate in each regions. Based on these parameters, the Asynchronous state was defined by changing empirical parameters in order to get not fluctuation in the brain regions. The result is a reduction of the spike-triggered adaptation of the excitatory neurons, a reduction of the number of connection between the regions, an augmentation of the inhibitory synaptic weights, a reduction of the variance of the noise and the addition of Poisson generator for the spiking neural network.

Based on the parameters of the Irregular Synchronous state, the parameters for the Regular Bursting state are chosen by changing the type of the neurons from regular spiking to regular burst neurons. This modification was done by changing the voltage reset of the membrane and the leak of the reversal potential of the excitatory and inhibitory neurons, the spike-triggered adaptation and time constant of the adaptation current of excitatory neurons. An empirical exploration of the models are done in order to get a balanced of the spiking neural network and the brain dynamic wanted. The result of this exploration is a reduction of the connection between regions and a reduction of the connection between excitatory and inhibitory neurons, a reduction of the number of connection between brain regions and a reduction of the variance of the noise. All the numerical values of the parameters are in the Supplementary Table 1.

### Communication between modules

#### Initialization of communication

During the initialization of the simulation, the launcher creates a specific folder for each module and an extra folder for the logger file of all components. The launcher creates a file with all the parameters. This file contains parameters for the simulation with dedicated sections for each module. The parameters shared between modules are duplicated by the launcher in each section to ensure there are the same.

#### Synchronization between modules

The transfer modules synchronize the simulation by managing the access to its internal buffer and receiving status messages from the simulators. The receiver process receives the data and aggregates them in a buffer. Rate data do not need to be buffered when use MPI communication, they are send or received directly to the transformer process. This buffer is transferred to the transformation function when the previous data are transformed and transferred to the sender process. The sender process gets the data after sending the previous data to the simulator. It can only send the data to the simulator when the simulator itself is ready. In addition, the simulator needs to await data for the next step of the simulation. The transfer module assures correct transport given all these constraints and keeps the components synchronized. If needed the transfer module buffers data for a simulation step. The transfer module can receive and send data concurrently and translation can be performed while waiting for the the slowest simulator.

### Performance test

The performance test is done by integrating recording of time at a specific place of the code. These times are aggregate duration to evaluate the running time of the co-simulation in each section. This allows evaluating the time of “simulation”, “IO” and “wait” time. Each test is done for 10 trails of a 1 biological second for asynchronous configurations with one or two parameters which vary per trial. Results of the trials are averaged to reduce the variability of the measurements. The varied parameters of the tests are the number of spiking neurons, synchronized time between simulators and the configuration of MPI and thread of NEST.

Figure 2 and supplementary figure **??, ??** and **??** show the result of the performance test done on DELL Precision-7540 (Intel Xeon(R) E-2286M CPU 2.40 GHz * 8 cores * 2 threads, 64 GB of Ram with Ubuntu 18.04.5). The communication between components in the transfer module was performed with the multithreading approach. Supplementary figure **??, ??** and **??** are generated using the Jusuf system (https://apps.fzjuelich.de/jsc/hps/jusuf/cluster/configuration.html) which is composed of nodes with 2 AMD EPYC 7742 2.25 GHz * 64 cores * 2 threads, 256 (16 16) GB DDR4 with 3200 MHz, connected by InfiniBand HDR100 (Connect-X6). In this second case, the transfer module uses MPI protocol for communication between components.

### Implementation details

The source code of the co-simulation is open-source and contains Python script and C++ files. A singularity and a docker image are also available on singularity-hubs for the replication of the figures as in the performance test. The activity diagram (see Supplementary Figure **??**) describes in detail the interaction between each module and components for this specific virtual experimentation.

The implementation details for the creation of the Input and Output for NEST is described in Supplementary Note 1. In addition to this note, an activity diagram (see Supplementary Figure **??**) describes the communication protocol with NEST back-end. For this specific example, the states of the wrapper of NEST and the states of transfer components which communicate with NEST are described respectively by the Supplementary Figure **??** and **??**.

In the same way, the description of the creation of the Input and Output for TVB is described in Supplementary Note 1. In addition to this note, an activity diagram (see Supplementary Figure **??**) describes the communication protocol with the TVB wrapper. For this specific example, the states of the wrapper of TVB and the states of transfer components which communicate with the wrapper of TVB are described respectively by the Supplementary Figures **??** and **??**.

The description of the transfer modules is partially described in the Supplementary Note 2 which focus only on the interface with simulators. In addition to this note, the state of the different components are described in the Supplementary Figure **??, ??** and **??**. For a better understanding between the different instances and classes of this module, the Supplementary Figure **??** describes all the instances and their role and the Supplementary Figure **??** describes the composition of the abstract class and the simple API for communication. The communication protocol for data exchange between component of the transfer modules are different depending on whether the parallelization strategy is multithreading or multiprocessing. In the case of multiprocessing, MPI protocol is used for data exchange. The communication protocol differs depending on the type of data as shown by the panel A of the Supplementary Figure **??**. The spike trains data are variable in size and rather large (more than 1 Megabit). The shared memory is chosen in this case. For the mean rate data, the size of data is rather constant and small (few Kilobits). Send and Receive function of MPI protocol is chosen in this case. In case of multi-threading, only shared buffer is used between thread.

### Dead lock due to the global interpreter of python

In the case of the transfer modules use multithreading for internal communication, it can happen that the program is in deadlock because the interface with a simulator does not receive a signal of a message. As it is explained on the global interpreter lock documentation, “The GIL(global interpreter lock) can cause I/O-bound threads to be scheduled ahead of CPU-bound threads, and it prevents signals from being delivered.” (https://wiki.python.org/moin/GlobalInterpreterLock). The consequence of it, that some signals used by MPI are not delivered, which create a situation where a simulator and a transformer are waiting a MPI message from the other one, this message will never arrive.

## References

[1] Finkelstein, A. et al. Computational challenges of systems biology. Com-puter 37 (5), 26–33 (2004). URL http://ieeexplore.ieee.org/document/1297236/. https://doi.org/10.1109/MC.2004.1297236.

[2] Einevoll, G. T. et al. The scientific case for brain simulations Neu-ron 102 (4), 735–744 (2019). URL https://hal.archives-ouvertes.fr/hal-02192934. https://doi.org/10.1016/j.neuron.2019.03.027, publisher: Elsevier.

[3] Meier-Schellersheim, M., Fraser, I. D. C. & Klauschen, F. Multiscale modeling for biologists. Wiley Interdiscip. Rev. Syst. Biol. Med. 1 (1), 4–14 (2009). URL http://onlinelibrary.wiley.com/doi/abs/10.1002/wsbm.33. https://doi.org/10.1002/wsbm.33.

[4] Schlick, T. et al. A MULTISCALE VISION-ILLUSTRATIVE APPLI-CATIONS FROM BIOLOGY TO ENGINEERING. Int. J. Mult. Comp. Eng. 19 (2), 39–73 (2021). URL http://www.dl.begellhouse.com/journals/61fd1b191cf7e96f,6cd7873d3aec47b1,75086009651d6035.html. https://doi.org/10.1615/IntJMultCompEng.2021039845.

[5] Rejniak, K. A. & Anderson, A. R. A. Hybrid models of tumor growth Wiley Interdiscip. Rev. Syst. Biol. Med. 3 (1), 115–125 (2011). URL https://www.ncbi.nlm.nih.gov/pmc/articles/PMC3057876/. https://doi.org/10.1002/wsbm.102.

[6] Rahman, M. M., Feng, Y., Yankeelov, T. E. & Oden, J. T. A fully coupled space–time multiscale modeling framework for predicting tumor growth Comput. Methods Appl. Mech. Eng. 320, 261–286 (2017). URL https://www.sciencedirect.com/science/article/pii/S004578251631249X. https://doi.org/10.1016/j.cma.2017.03.021.

[7] Durstewitz, D., Seamans, J. K. & Sejnowski, T. J. Neurocomputational models of working memory Nature Neurosci. 3 (11), 1184–1191 (2000). URL http://www.nature.com/articles/nn11001184. https://doi.org/10.1038/81460, xnumber: 11 Publisher: Nature Publishing Group.

[8] COMSOL multiphysics reference manual.

[9] MATLAB. version 9.2.0 (R2017a) (The MathWorks Inc., Natick, Massachusetts, 2017).

[10] Fish, J., Wagner, G. J. & Keten, S. Mesoscopic and multiscale modelling in materials. Nat. Mater. 20 (6), 774–786 (2021). URL http://www.nature.com/articles/s41563-020-00913-0. https://doi.org/10.1038/s41563-020-00913-0, number: 6 Primary atype: Reviews Pub-lisher: Nature Publishing Group Subject term: Engineering;Materials sci-ence;Mathematics and computing Subject term id: engineering;materials-science;mathematics-and-computing.

[11] Gomes, C., Thule, C., Broman, D., Larsen, P. G. & Vangheluwe, H. Co-simulation: A survey. ACM Comput. Surv. 51 (3), 1–33 (2018). URL https://doi.org/10.1145/3179993.

[12] Matthews, M. L. & Marshall-Colón, A. Multiscale plant modeling: from genome to phenome and beyond Emerg. Top. Life Sci. 5 (2), 231–237 (2021). URL https://doi.org/10.1042/ETLS20200276. https://doi.org/10.1042/ETLS20200276.

[13] Hetherington, J. et al. Addressing the challenges of multiscale model management in systems biology. Comput. Chem. Eng. 31 (8), 962–979 (2007). URL https://www.sciencedirect.com/science/article/pii/S0098135406002535. https://doi.org/10.1016/j.compchemeng.2006.10.004.

[14] Carnevale, N. T. & Hines, M. L. The NEURON Book Cambridge Uni-versity Press, (2006). URL https://www.cambridge.org/core/books/neuron-book/7C8D9BD861D288E658BEB652F593F273.

[15] Akar, N. A. et al. Leporati, F., Danese, G., Torti, E. & D’Agostino, D. (eds) Arbor — a morphologically-detailed neural network simulation library for contemporary high-performance computing architectures. (eds Leporati, F., Danese, G., Torti, E. & D’Agostino, D.) 2019 27th Euromi-cro International Conference on Parallel, Distributed and Network-Based Processing (PDP), 274–282 (2019).

[16] Bower, J. M. & Beeman, D. in Introduction (eds Bower, J. M. & Beeman, D.) The Book of GENESIS: Exploring Realistic Neural Models with the GEneral NEural SImulation System 3–5 (Springer, 1998). URL https://doi.org/10.1007/978-1-4612-1634-61.

[17] Gewaltig, M.-O. & Diesmann, M. NEST (NEural simulation tool). Schol-arpedia 2 (4), 1430 (2007). URL http://www.scholarpedia.org/article/NEST(NEuralSimulation_Tool). https://doi.org/10.4249/scholarpedia.1430.

[18] Stimberg, M., Brette, R. & Goodman, D. F. Brian 2, an intuitive and efficient neural simulator. eLife 8, e47314 (2019).https://doi.org/10.7554/eLife.47314.

[19] Sanz Leon, P. et al. The virtual brain: a simulator of primate brain network dynamics. Front. Neuroinform. 7 (2013). URL https://www.frontiersin.org/articles/10.3389/fninf.2013.00010/full. https://doi.org/10.3389/fninf.2013.00010.

[20] Cakan, C., Jajcay, N. & Obermayer, K. neurolib: A simulation framework for whole-brain neural mass modeling. Cogn. Comput. (2021). URL https://doi.org/10.1007/s12559-021-09931-9SMASH. https://doi.org/10.1007/s12559-021-09931-9SMASH.

[21] Goddard, N., Hood, G., Howell, F., Hines, M. & De Schutter, E. NEOSIM: Portable large-scale plug and play modelling Neurocomputing 38-40, 1657–1661 (2001). URL https://www.sciencedirect.com/science/article/pii/S0925231201005288. https://doi.org/10.1016/S0925-2312(01)00528-8

[22] Tony, P. BRAHMS: Novel middleware for integrated systems compu-tation Adv. Eng. Inform. 24 (1), 49–61 (2010). URL http://www.frontiersin.org/10.3389/conf.neuro.11.2008.01.051/eventabstract. https://doi.org/10.3389/conf.neuro.11.2008.01.051.

[23] Falotico, E. et al. Connecting artificial brains to robots in a compre-hensive simulation framework: The neurorobotics platform Frontiers in Neurorobotics 11, 2 (2017). URLhttps://www.frontiersin.org/article/10.3389/fnbot.2017.00002.

[24] Djurfeldt, M. et al. Run-time interoperability between neuronal network simulators based on the MUSIC framework. Neuroinformatics 8 (1), 43–60 (2010). URL https://doi.org/10.1007/s12021-010-9064-z. https://doi.org/10.1007/s12021-010-9064-z.

[25] Renz Aline F., Jihyun Lee, Klas Tybrandt, Maciej Brzezinski, Dayra Lorenzo, Mouna Cerra Cheraka, Jaehong Lee, Fritjof Helmchen, Janos Vörös, and Christopher M. Lewis. “Opto-E-Dura: A Soft, Stretchable ECoG Array for Multimodal, Multi-Scale Neuroscience.” BioRxiv, July 1, 2020, 2020.06.10.139493. https://doi.org/10.1101/2020.06.10.139493. Renz, A. F., Lee, J., Tybrandt, K., Brzezinski, M., Klas, T., Cerra Cheraka, M., Jaehong, J., Helmchen, F., Vörös, J., Lewis, M. C. Opto-e-dura: A soft, stretchable ECoG array for multimodal, multiscale neuroscience. Advanced Healthcare Materials 9 (17), 2000814 (2020). URL http://onlinelibrary.wiley.com/doi/abs/10.1002/adhm.202000814. https://doi.org/https://doi.org/10.1002/adhm.202000814, eprint: https://onlinelibrary.wiley.com/doi/pdf/10.1002/adhm.202000814.

[26] Lein, E. S. et al. Genome-wide atlas of gene expression in the adult mouse brain. Nature 445 (7124), 168–176 (2007). URL http://www.nature.com/articles/nature05453. https://doi.org/10.1038/nature05453.

[27] Hahne, J. et al. NEST 3.0 (2021). URL https://zenodo.org/record/4739103.

[28] Shuman, T. et al. Breakdown of spatial coding and interneuron syn-chronization in epileptic mice. Nat. Neurosci. 23 (2), 229–238 (2020). URL https://www.ncbi.nlm.nih.gov/pmc/articles/PMC7259114/. https://doi.org/10.1038/s41593-019-0559-0.

[29] Oh, S. W. et al. A mesoscale connectome of the mouse brain. Nature 508 (7495), 207–214 (2014). URL https://www.nature.com/articles/nature13186. https://doi.org/10.1038/nature13186.

[30] di Volo, M., Romagnoni, A., Capone, C. & Destexhe, A. Biologically real-istic mean-field models of conductance-based networks of spiking neurons with adaptation. Neural Computation 31 (4), 653–680 (2019). URL https://www.mitpressjournals.org/doi/10.1162/necoa01173. https://doi.org/10.1162/necoa01173.

[31] Brette, R. & Gerstner, W. Adaptive exponential integrate-and-fire model as an effective description of neuronal activity. Journal of Neurophysiology 94 (5), 3637–3642 (2005). URL http://www.physiology.org/doi/10.1152/jn.00686.2005. https://doi.org/10.1152/jn.00686.2005.

[32] Sanz-Leon, P., Knock, S. A., Spiegler, A. & Jirsa, V. K. Mathemat-ical framework for large-scale brain network modeling in the virtual brain. NeuroImage 111, 385–430 (2015). URL http://www.sciencedirect.com/science/article/pii/S1053811915000051. https://doi.org/10.1016/j.neuroimage.2015.01.002.

[33] Hagen, E. et al. Hybrid scheme for modeling local field potentials from point-neuron networks. Cereb. Cortex 26 (12), 4461–4496 (2016). URL https://academic.oup.com/cercor/article/26/12/4461/2333943. https://doi.org/10.1093/cercor/bhw237.

[34] Kuhn, A., Aertsen, A. & Rotter, S. Higher-order statistics of input ensembles and the response of simple model neurons. Neural Computation 15 (1), 67–101 (2003). URL http://www.mitpressjournals.org/doi/10.1162/089976603321043702. https://doi.org/10.1162/089976603321043702.

[35] Chopard, B., Borgdorff, J. & Hoekstra, A. G. A framework for multi-scale modelling. Philosophical Transactions of the Royal Society A: Mathe-matical, Physical and Engineering Sciences 372 (2021), 20130378 (2014). URL https://royalsocietypublishing.org/doi/full/10.1098/rsta.2013.0378. https://doi.org/10.1098/rsta.2013.0378, publisher: Royal Society.

[36] IEEE standard for modeling and simulation (M & S) high level architecture (HLA)– framework and rules. IEEE Std 1516-2010 1–38 (2010).https://doi.org/10.1109/IEEESTD.2010.5553440, conference Name: IEEE Std 1516-2010 (Revision of IEEE Std 1516-2000).

[37] Andreas Junghanns et al. Sjölund, M., Buffoni, L., Pop, A. & Ochel, L. (eds) The functional mock-up interface 3.0 -new features enabling new applications. (eds Sjölund, M., Buffoni, L., Pop, A. & Ochel, L.) Proceedings of 14th Modelica Conference 2021, Linköping, Sweden, September 20-24, 2021, 17–26 (2021). URL https://ecp.ep.liu.se/index.php/modelica/article/view/178.

[38] Boustani, S. E. & Destexhe, A. A master equation formalism for macroscopic modeling of asynchronous irregular activity states. Neural computation21 46–100 (2009).

[39] Coveney, P. V., Groen, D. & Hoekstra, A. G. Reliability and repro-ducibility in computational science: implementing validation, verification and uncertainty quantification in silico. Philosophical Transactions of the Royal Society A: Mathematical, Physical and Engineering Sci-ences 379 (2197), 20200409 (2021). URLhttps://royalsocietypublishing.org/doi/10.1098/rsta.2020.0409. https://doi.org/10.1098/rsta.2020.0409, publisher: Royal Society.

[40] Coveney, P. V. & Highfield, R. R. When we can trust computers (and when we can’t). Philosophical Transactions of the Royal Society A: Mathematical, Physical and Engineering Sciences 379 (2197), 20200067 (2021). URL https://royalsocietypublishing.org/doi/10.1098/rsta.2020.0067. https://doi.org/10.1098/rsta.2020.0067, publisher: Royal Society.

[41] Taveres-Cachat, E., Favoino, F., Loonen, R. & Goia, F. Ten questions concerning co-simulation for performance prediction of advanced building envelopes. Building and Environment 191, 107570 (2021). URL https://www.sciencedirect.com/science/article/pii/S0360132320309379. https://doi.org/10.1016/j.buildenv.2020.107570.

[42] Belaud, J.-P. & Pons, M. in Open software architecture for process simu-lation: The current status of CAPE-OPEN standard (eds Grievink, J. & van Schijndel, J.) Computer Aided Chemical Engineering, Vol. 10 of Euro-pean Symposium on Computer Aided Process Engineering-12 847–852 (Elsevier, 2002). URL https://www.sciencedirect.com/science/article/pii/S1570794602801699.

[43] Braun, R. & Krus, P. Multi-threaded distributed system simulations using the transmission line element method. SIMULATION 92 (10), 921–930 (2016). URL https://doi.org/10.1177/0037549716667243. https://doi.org/10.1177/0037549716667243, publisher: SAGE Publications Ltd STM.

[44] Nguyen, V. H., Besanger, Y., Tran, Q. T. & Nguyen, T. L. On con-ceptual structuration and coupling methods of co-simulation frameworks in cyber-physical energy system validation. Energies 10 (12), 1977 (2007). URL https://www.mdpi.com/1996-1073/10/12/1977. https://doi.org/10.3390/en10121977, xnumber: 12 publisher: Multidisciplinary Digital Publishing Institute.

[45] Nordlie, E., Gewaltig, M.-O. & Plesser, H. E. Towards repro-ducible descriptions of neuronal network models. PLOS Computa-tional Biology 5 (8), e1000456 (2009). URL https://journals.plos.org/ploscompbiol/article?id=10.1371/journal.pcbi.1000456. https://doi.org/10.1371/journal.pcbi.1000456, publisher: Public Library of Science.

[46] Melozzi, F., Woodman, M. M., Jirsa, V. K. & Bernard, C. The vir-tual mouse brain: A computational neuroinformatics platform to study whole mouse brain dynamics. eNeuro 4 (3), ENEURO.0111–17.2017 (2017). URLhttp://www.eneuro.org/content/4/3/ENEURO.0111-17.2017. https://doi.org/10.1523/ENEURO.0111-17.2017.

## References

[1] Nordlie, E., Gewaltig, M.-O. & Plesser, H. E. Towards reproducible descriptions of neuronal network models. PLOS Computational Biology 5 (8), e1000456 (2009). URLhttps://journals.plos.org/ploscompbiol/article?id=10.1371/journal.pcbi.1000456. https://doi.org/10.1371/journal.pcbi.1000456, publisher: Public Library of Science.

[2] Hahne, J. et al. NEST 3.0 (2021). URLhttps://zenodo.org/record/4739103.

[3] Shuman, T. et al. Breakdown of spatial coding and interneuron synchronization in epileptic mice. Nat. Neurosci. 23 (2), 229–238 (2020). URL https://www.ncbi.nlm.nih.gov/pmc/articles/PMC7259114/. https://doi.org/10.1038/s41593-019-0559-0.

[4] Brette, R. & Gerstner, W. Adaptive exponential integrate-and-fire model as an effective description of neuronal activity. Journal of Neurophysiology 94 (5), 3637–3642 (2005). URL http://www.physiology.org/doi/10.1152/jn.00686.2005. https://doi.org/10.1152/jn.00686.2005.

[5] Sanz-Leon, P., Knock, S. A., Spiegler, A. & Jirsa, V. K. Mathematical framework for large-scale brain network modeling in the virtual brain. NeuroImage 111, 385–430 (2015). URL http://www.sciencedirect.com/science/article/pii/S1053811915000051. https://doi.org/10.1016/j.neuroimage.2015.01.002.

[6] Oh, S. W. et al. A mesoscale connectome of the mouse brain. Nature 508 (7495), 207–214 (2014). URL https://www.nature.com/articles/nature13186. https://doi.org/10.1038/nature13186.

[7] di Volo, M., Romagnoni, A., Capone, C. & Destexhe, A. Biologically realistic mean-field models of conductance-based networks of spiking neurons with adaptation. Neural Computation 31 (4), 653–680 (2019). URL https://www.mitpressjournals.org/doi/10.1162/necoa01173. https://doi.org/10.1162/necoa01173.

[8] Sanz-Leon, P., Knock, S. A., Spiegler, A. & Jirsa, V. K. Mathematical framework for large-scale brain network modeling in the virtual brain. NeuroImage 111, 385–430 (2015). URL http://www.sciencedirect.com/science/article/pii/S1053811915000051. https://doi.org/10.1016/j.neuroimage.2015.01.002.

[9] Hagen, E. et al. Hybrid scheme for modeling local field potentials from point-neuron networks. Cereb. Cortex 26 (12), 4461–4496 (2016). URL https://academic.oup.com/cercor/article/26/12/4461/2333943. https://doi.org/10.1093/cercor/bhw237.

