## Supplementary figure for "Multiscale cosimulation design template for neuroscience applications"

| Supplementary File | Title |
| --- | --- |
| Supplementary Figure 1 | Zoom on 1s for the spiking neural network with regular bursting state |
| Supplementary Figure 2 | Detail sequence diagram of the co-simulation |
| Supplementary Figure 3 | Sequence diagram of the communication protocol with NEST |
| Supplementary Figure 4 | State diagram of NEST wrapper |
| Supplementary Figure 5 | State diagram of transfer components for interaction with NEST |
| Supplementary Figure 6 | Sequence diagram of the communication protocol with the wrapper of TVB |
| Supplementary Figure 7 | State diagram of wrapper of TVB modules |
| Supplementary Figure 8 | State diagram of transfer components for interaction with the wrapper of TVB |
| Supplementary Figure 9 | File organisation of the transformer module |
| Supplementary Figure 10 | State diagram of transfer components which transform data from one scale to another |
| Supplementary Figure 11 | Structure diagram of the two transfer modules with the description of each component |
| Supplementary Figure 12 | Class diagram of the transfer modules |
| Supplementary Figure 13 | Internal communication in the transfer modules between the components |
| Supplementary Figure 14 | Performance of the co-simulation on one computer with different number of neurons |
| Supplementary Figure 15 | Performance of the co-simulation on one computer with different time of synchronization between simulators |
| Supplementary Figure 16 | Performance of the co-simulation on one computer with different process and thread for NEST |
| Supplementary Figure 17 | Performance of the co-simulation on one supercomputer for different number of neurons |
| Supplementary Figure 18 | Performance of the co-simulation on one supercomputer for different time of synchronization between simulator |
| Supplementary Figure 19 | Performance of the co-simulation on one supercomputer for different number of node for NEST |
| Supplementary Figure 20 | Details of the timer of one run for the reference configuration |
| Supplementary Figure 21 | Proof of concept of replacing NEST and TVB by other simulators |
| Supplementary Tabular 1 | Tabular describing the co-simulation |
| Supplementary Note 1 | Guidelines Input/Output (I/O) interface |
| Supplementary Note 2 | Guideline for implementation of transfer module |
| Supplementary Note 3 | Detail characterization of the workflow TVB-NEST |

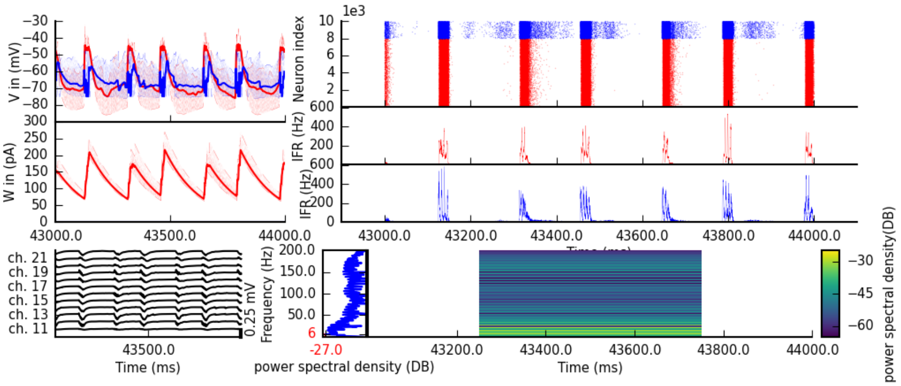


**Supplementary Figure 1** Zoom on 1s for the spiking neural network with regular bursting state

This figure is a zoom of the figure 3 between 43s and 44s. **top-left** Example of time series from 10 adaptive exponential leaky and integrator neurons. The red lines are excitatory and the blue curve are inhibitory neurons. The mean excitatory time series is shown with a thick red line and the inhibitory time series is shown with a thick blue line. **middle-left** The adaptation currents of 10 neurons are shown. The thick line is the mean adaptive currents. **bottom-left** The figure shown local field potential from the 12 sites of in the middle line of the polytrode. The local field potential is computed from the spike trains of all neurons by the software HybridLFPY[4]. **top-right** The figure shown spike trains of 10000 neurons for 1s. **bottom-right** The figure shown respectively the excitatory and inhibitory instantaneous firing rate of the population in panel middle-right in red and blue. **bottom** Spectrogram and power spectrum example of the instantaneous firing rate for 1s.

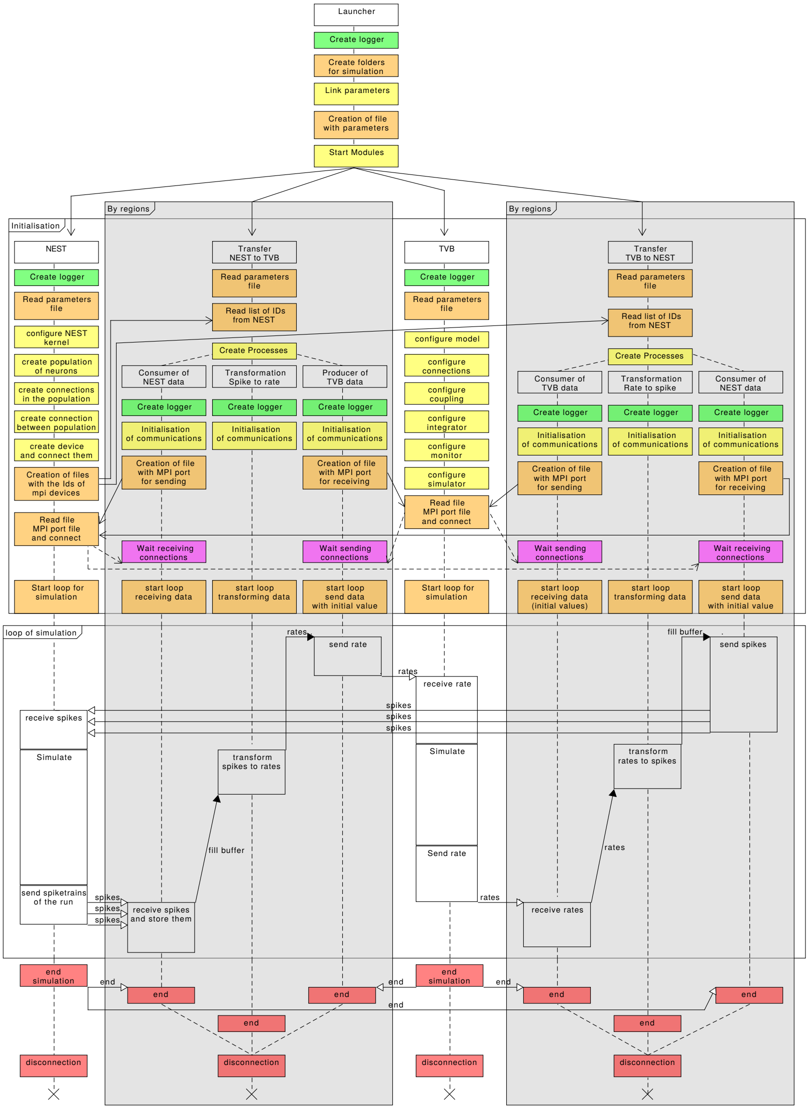

**Supplementary Figure 2** Detail sequence diagram of the co-simulation
This figure represents the interaction among the different modules during the co-simulation and the different exchanged data. The co-simulation is separated in 3 steps: initialisation and configuration, simulation and termination.
The colour code of the boxes :

- green for the creation of a logger, one by components and modules
- orange for access to file systems (the creation of a folder or a file, the reading of files, ...) and the start of the simulation with initial condition.
- yellow for initialization and configuration of modules and components
- magenta for MPI waiting connections
- white for the simulation step and the name of modules or components
- red for the termination of the simulation.

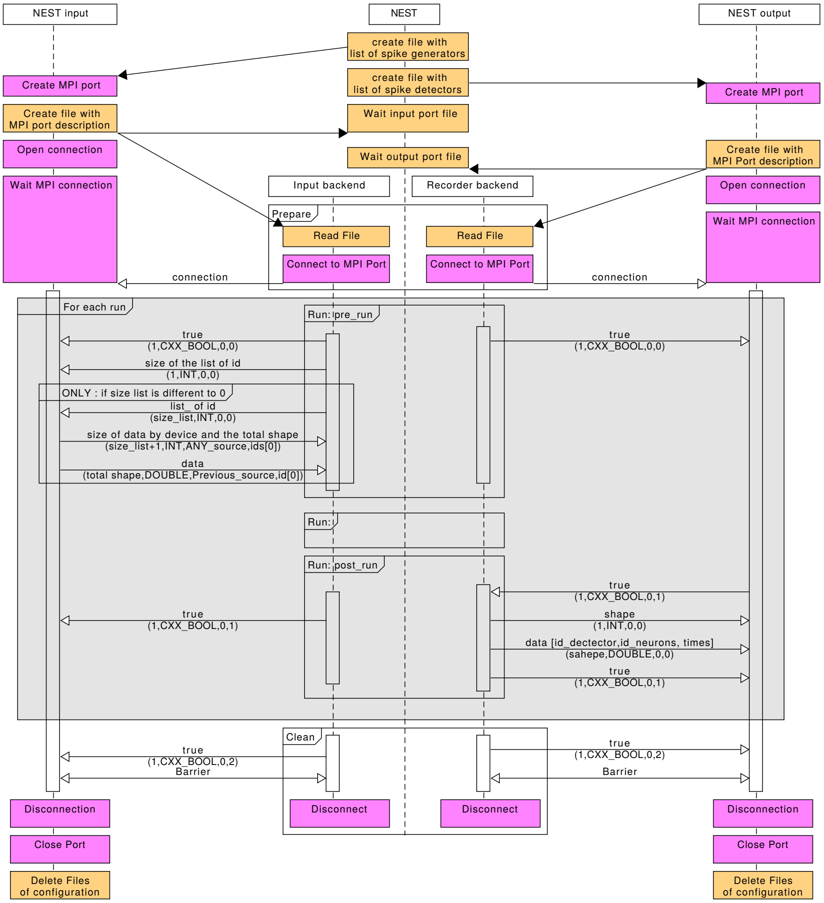


**Supplementary Figure 3** Sequence diagram of the communication protocol with NEST
The communication with NEST[1] is separated in 3 steps: creation of the MPI connection, simulation and termination.
The colour code of the boxes :

- orange for access to file systems (the creation of files, the reading of files, ...).
- magenta for management of MPI port
- white for the name of the modules or components

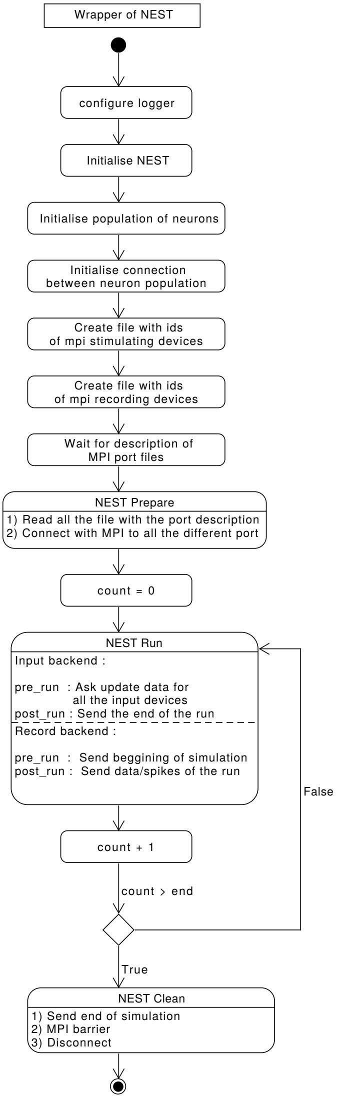


**Supplementary Figure 4** State diagram of NEST wrapper
The diagram describe all the different state of the NEST wrapper during the co-simulation. The beginning is the set-up of the network (the creation of neurons, their connection and the creation of devices). After, the loop of simulation and the termination.

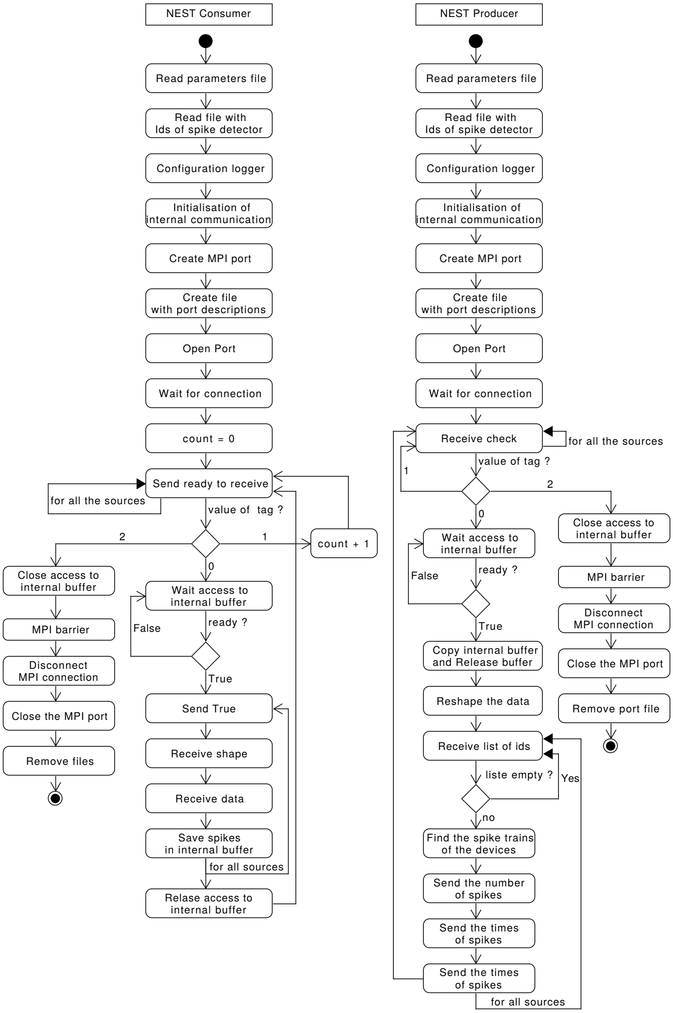


**Supplementary Figure 5** State diagram of transfer components for interaction with NEST

The diagram describes all the state of the component of the transfer module which communicates with NEST. The beginning is the configuration of itself and the creation of the MPI connection. Once the MPI connection is made, there is the loop of the simulation. The centre of the simulation loop is the value of the tag receive by the component to identify if NEST is ready to receive or send messages. If this tag equals 2, the components go in the sequence for the termination phase.

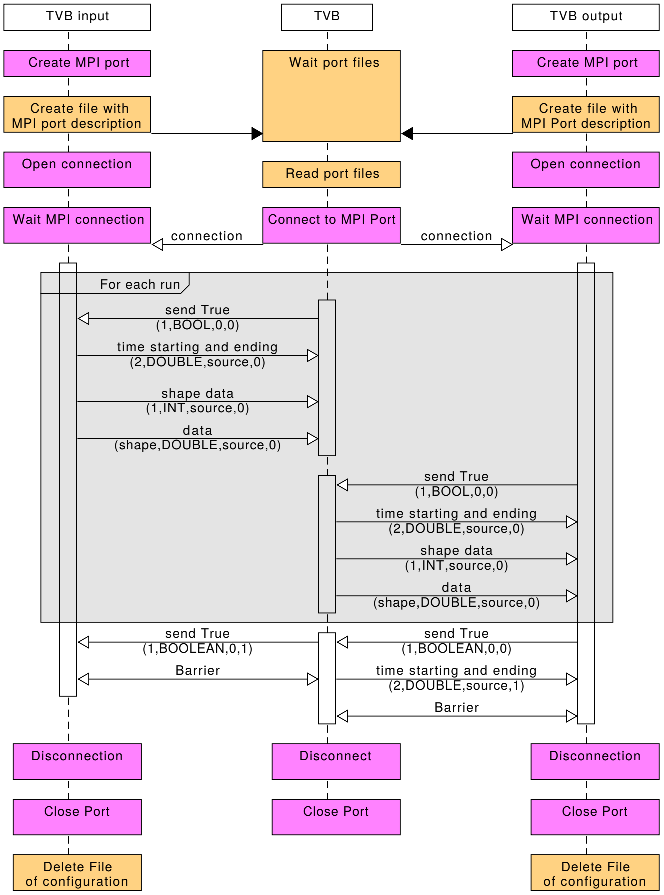


**Supplementary Figure 6** Sequence diagram of the communication protocol with the wrapper of TVB.

The communication with TVB[2] is separated in 3 steps: creation of the MPI connection, simulation and termination.

The colour code of the boxes :

- orange for access to file systems (create files, read files, ...).
- magenta for management of MPI port
- white for the name of the modules or components

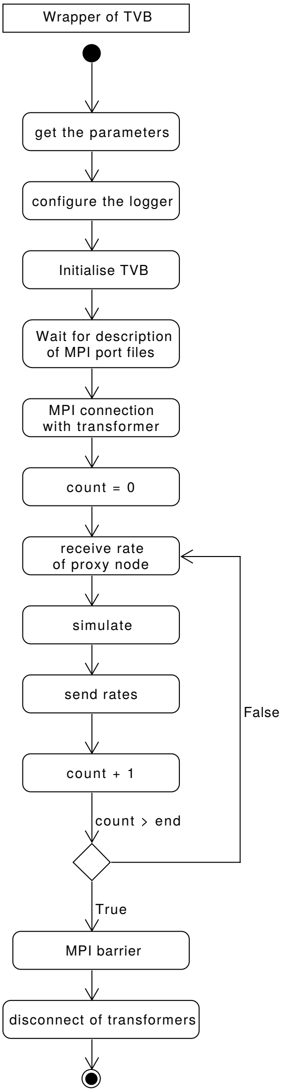


**Supplementary Figure 7** State diagram of wrapper of TVB modules
The diagram describe all the different state of the TVB wrapper during the co-simulation. The beginning is the set-up of the network (the creation of neurons, their connection and the creation of devices). After, the loop of simulation and the termination.

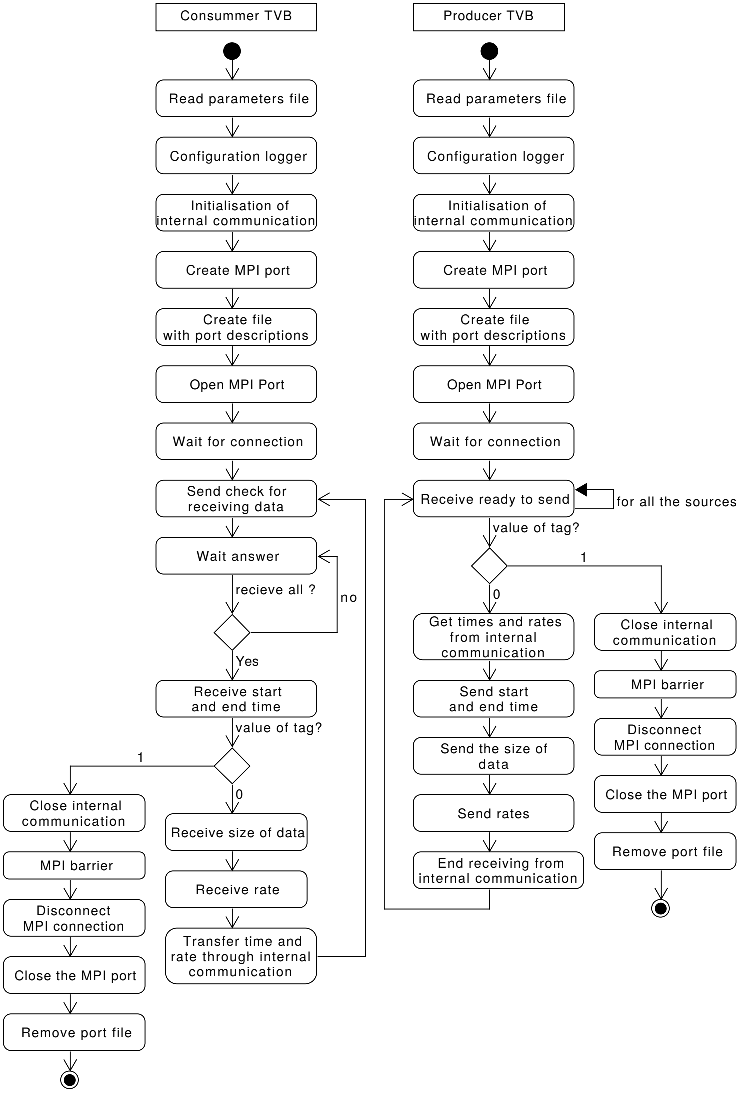


**Supplementary Figure 8** State diagram of transfer components for interaction with the wrapper of TVB

The diagram describes all the state of the component of the transfer module which communicates with TVB. The beginning is the configuration of itself and the creation of the MPI connection. Once the MPI connection is made, there is the loop of the simulation. The centre of the simulation loop is the value of the tag receive by the component to identify if NEST is ready to receive or send messages. If this tag equals 1, the components go in the sequence for the termination phase.

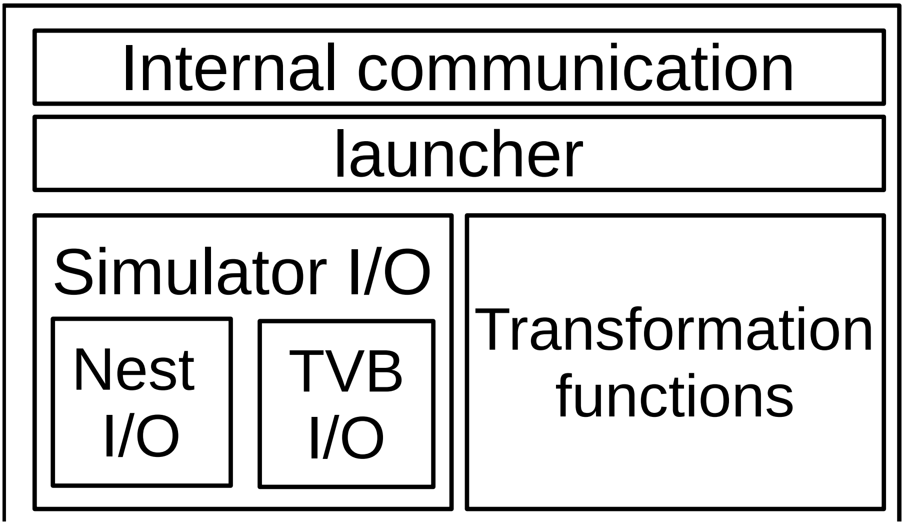


**Supplementary Figure 9** File organisation of the transformer module
The file of transfer modules are organised following the modularity of the module. The folder internal communication contains all the functions of the communication between components of the modules. The launcher regroups the files used to start the modules of transformation between specific simulators. The Simulator I/O contains the function for the interface of each simulator. The figure shows that interface for each simulator (TVB and NEST) is separated and independent. Transformation functions is a folder which contains the abstract class for the transformations and the implementation of specific transformation.

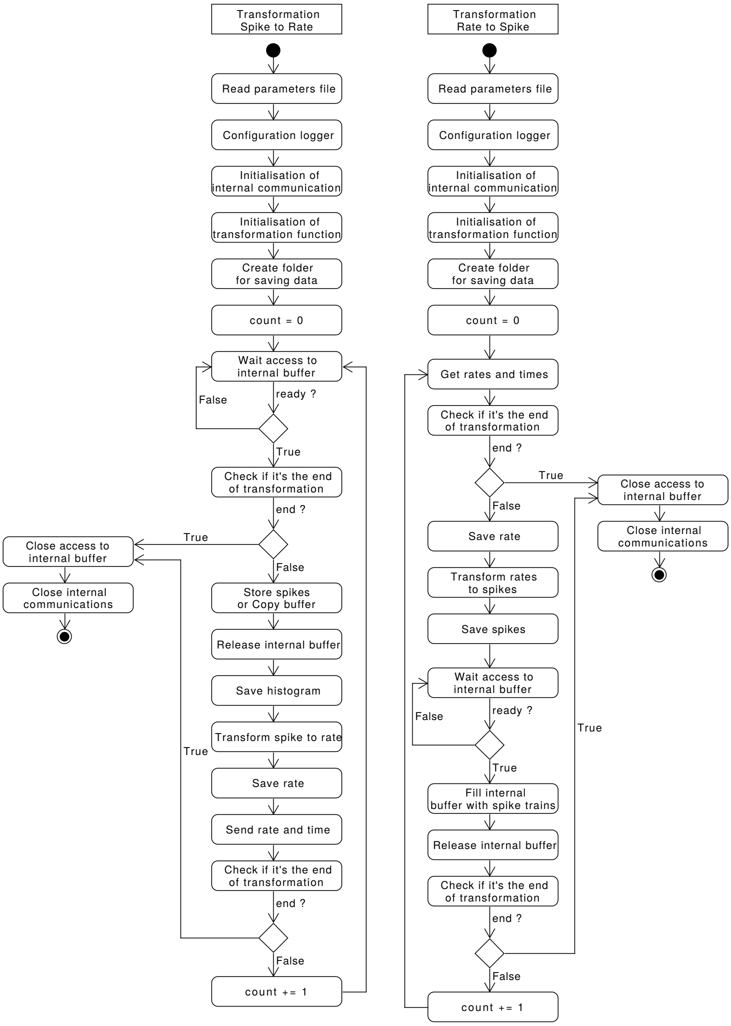


**Supplementary Figure 10** State diagram of transfer components which transform data from one scale to another

The diagram describes all the state of the transfer components. The beginning is the configuration of itself. The component is waiting to access to the data from one buffer for the transformation. After accessing to the data, it transforms them and awaits the access for writing in a second buffer. When it receives the termination from one side or the other, it goes in the sequence of termination.

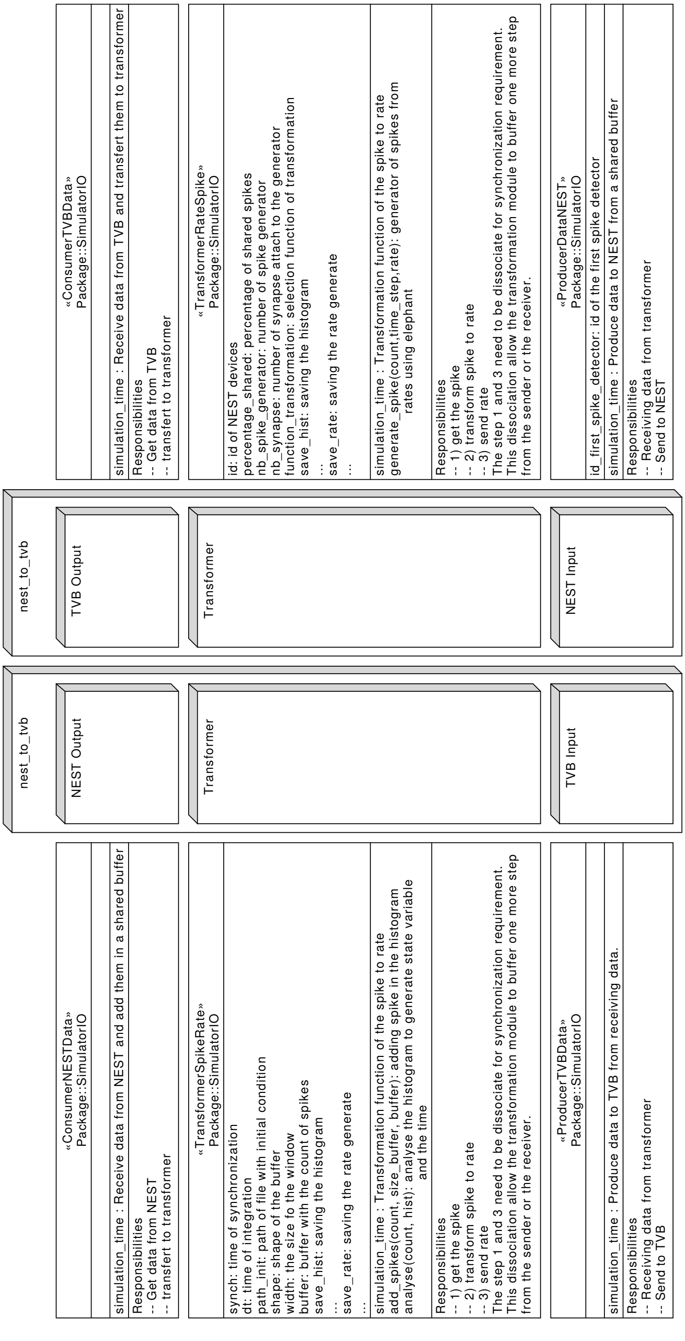


**Supplementary Figure 11** Structure diagram of the two transfer modules with the description of each component

Each component is based on a class. The diagram gives a short description of each class and their contents.

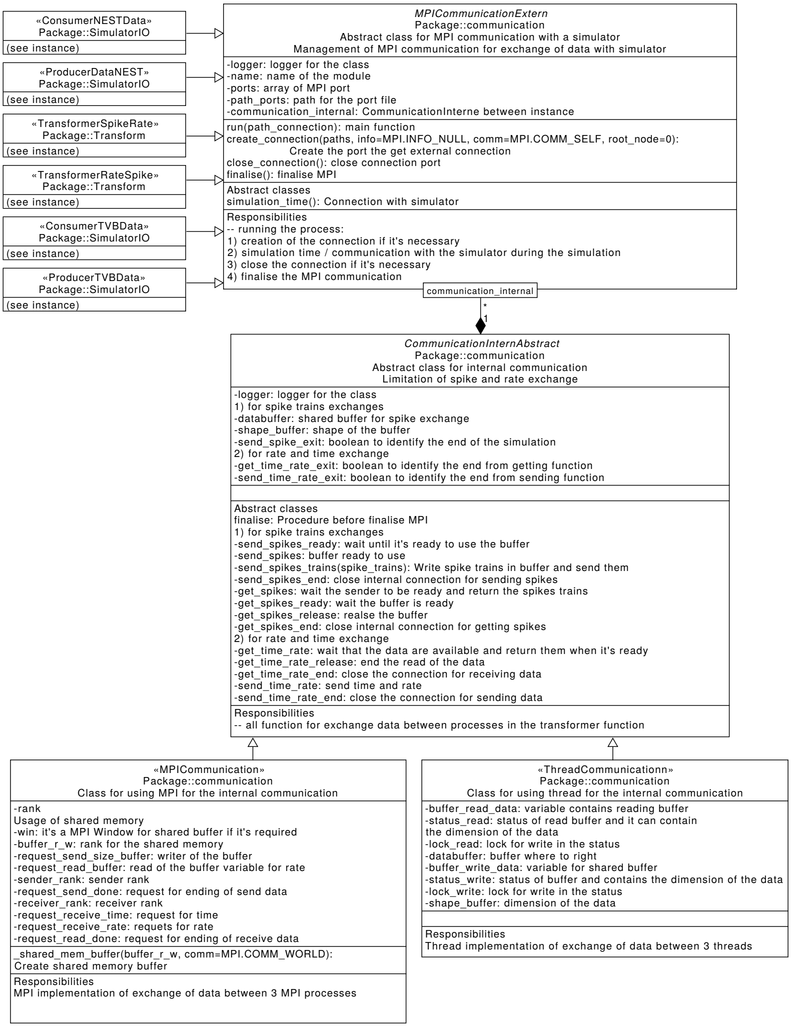


**Supplementary Figure 12** Class diagram of the transfer modules.

The description of the class on the right part is described in the supplementary figure 11. All this class inherit from an abstract class which manage MPI communication. This abstract class manage the MPI connection and has an internal communicator. This internal communicator is an abstract class for the communication between the transfer components. This internal communication can be melting process or multithreading as the diagram shows.

a

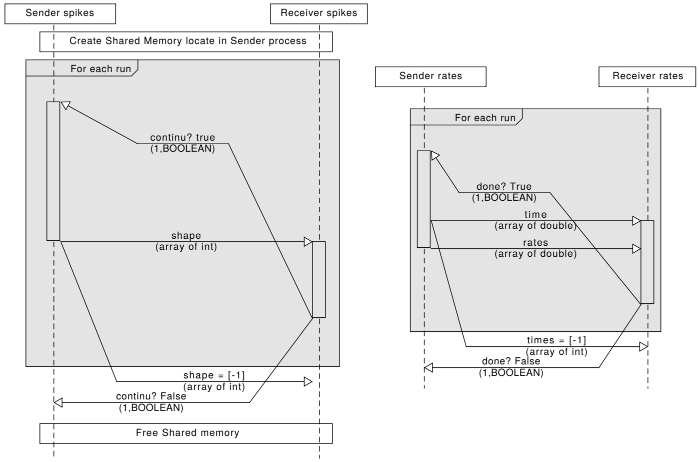


b

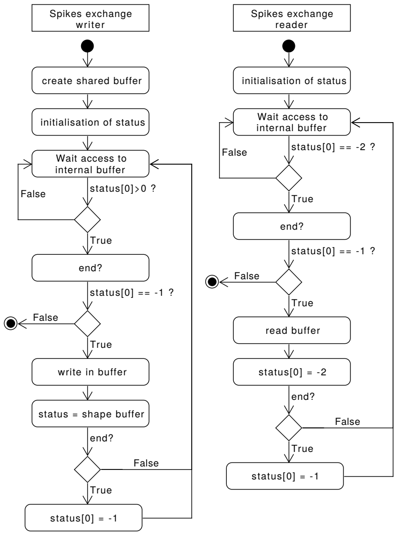


**Supplementary Figure 13** Communication between components in the transfer module
As described in the supplementary figure 12, the internal communication has 2 implementations. The panel a is a sequence diagram of the communication of spikes and rates using MPI communication. The panel b is a state diagram of the management of a shared buffer in the case of multithreading communication for transfer spike data.

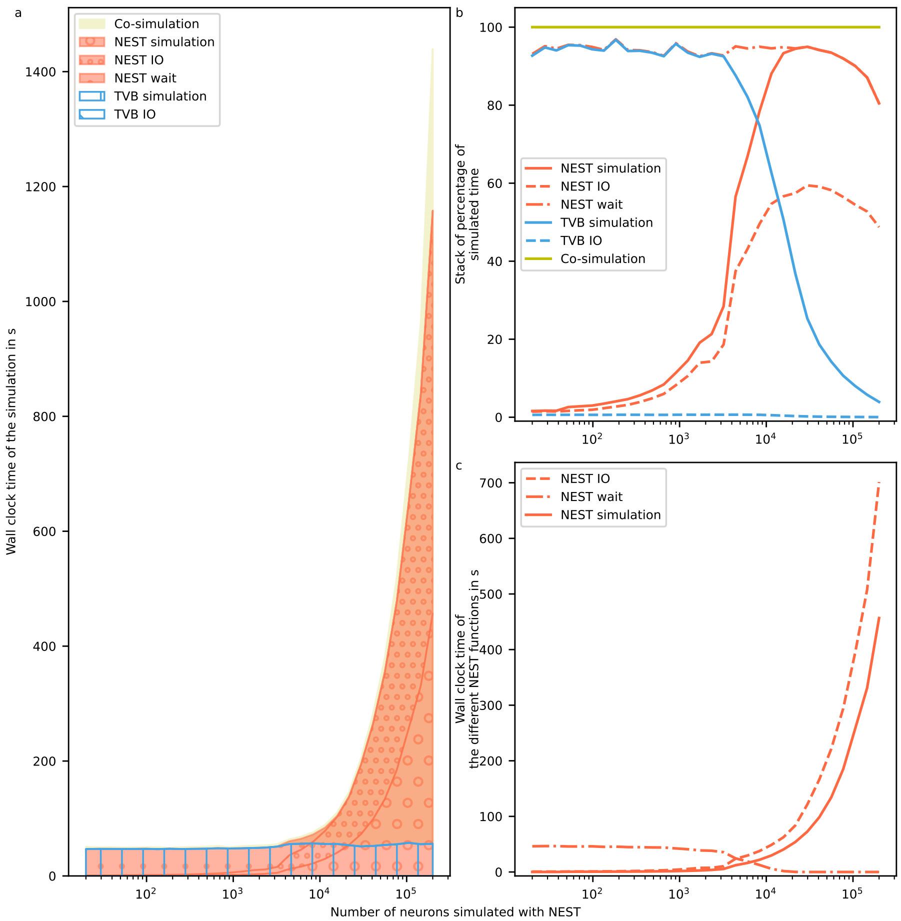


**Supplementary Figure 14** Details of the performance with the increase of neurons Performance is obtained for 1 biological second on a computer (see Online Method for more details). The reference implementation use 1 MPI process, 6 virtual processes/threads, 2.0 ms to synchronize time between simulator for the simulation of 20000 neurons. Simulation time depending on the number of neurons simulated with NEST. **a** The wall clock time of the simulator depending on the number of neurons. The total time of the co-simulation is represented in yellow. The "simulation", "IO" and "wait" times of NEST are represented in red surface with respectively hatches with big circles, small circles and points. The "simulation" and "IO" times of TVB are represented in the blue surface with respectively hatches horizontal lines and oblique lines. **b** The wall clock time for the co-simulation (yellow curve), NEST (red curves) and TVB (blue curves) by the total wall clock time. The solid, dashed and dashed dotted curves are associated with "simulation", "IO" and "wait" time of NEST. The solid and dashed line is associate to "simulation" and "IO" time of TVB. **c** The different timer for NEST simulator. Each contribution is reported as red curve and for increasing numbers of neurons. The solid, dashed and dashed dotted curves represent "simulation", "IO" and "wait" time of NEST respectively.

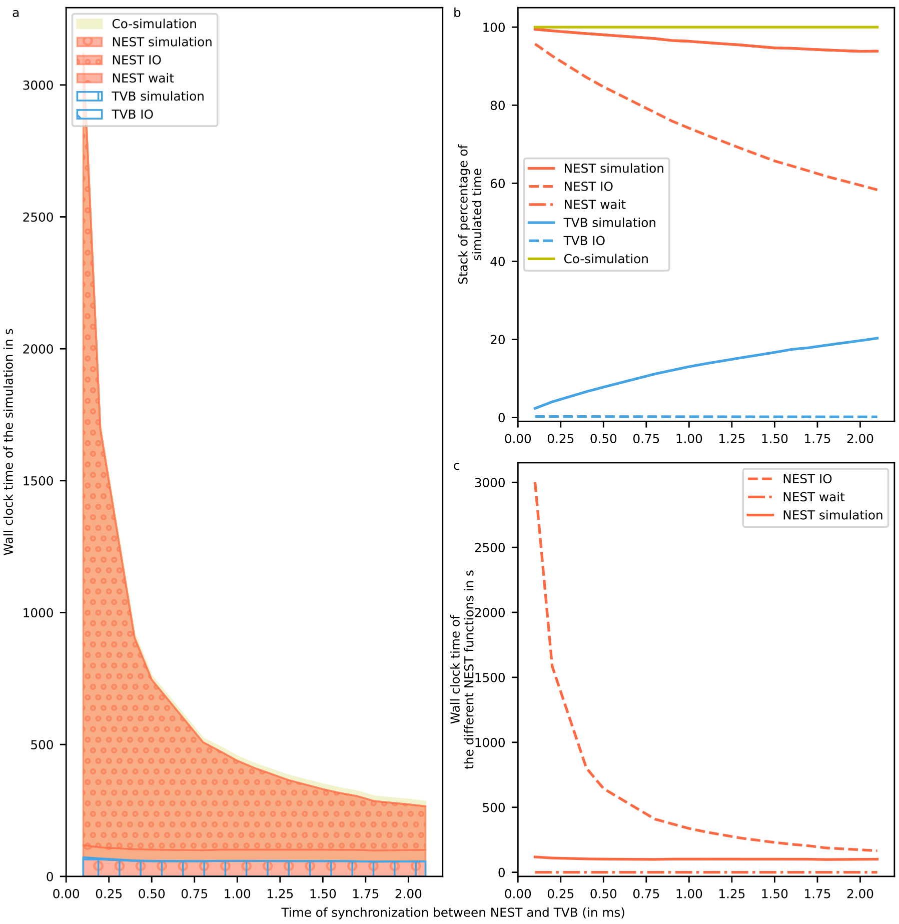


**Supplementary Figure 15** Details of the performance with the increase of synchronize time

Performance is obtained for 1 biological second on a computer (see Online Method for more details). The reference implementation use 1 MPI process, 6 virtual processes/threads, 2.0 ms to synchronize time between simulator for the simulation of 20000 neurons. Simulation time depending on the synchronized time between NEST and TVB. **a** The wall clock time of the simulator depending on time of synchronization between the two simulators. The total time of the co-simulation is represented in yellow. The "simulation", "IO" and "wait" times of NEST are represented in red surface with respectively hatches with big circles, small circles and points. The "simulation" and "IO" times of TVB are represented in the blue surface with respectively hatches horizontal lines and oblique lines. **b** The wall clock time for the co-simulation (yellow curve), NEST (red curves) and TVB (blue curves) by the total wall clock time. The solid, dashed and dashed dotted curves are associated with "simulation", "IO" and "wait" time of NEST. The solid and dashed line is associate to "simulation" and "IO" time of TVB. **c** The different timer for NEST simulator. Each contribution is reported as red curve and for increasing numbers of neurons. The solid, dashed and dashed dotted curves represent "simulation", "IO" and "wait" time of NEST respectively.

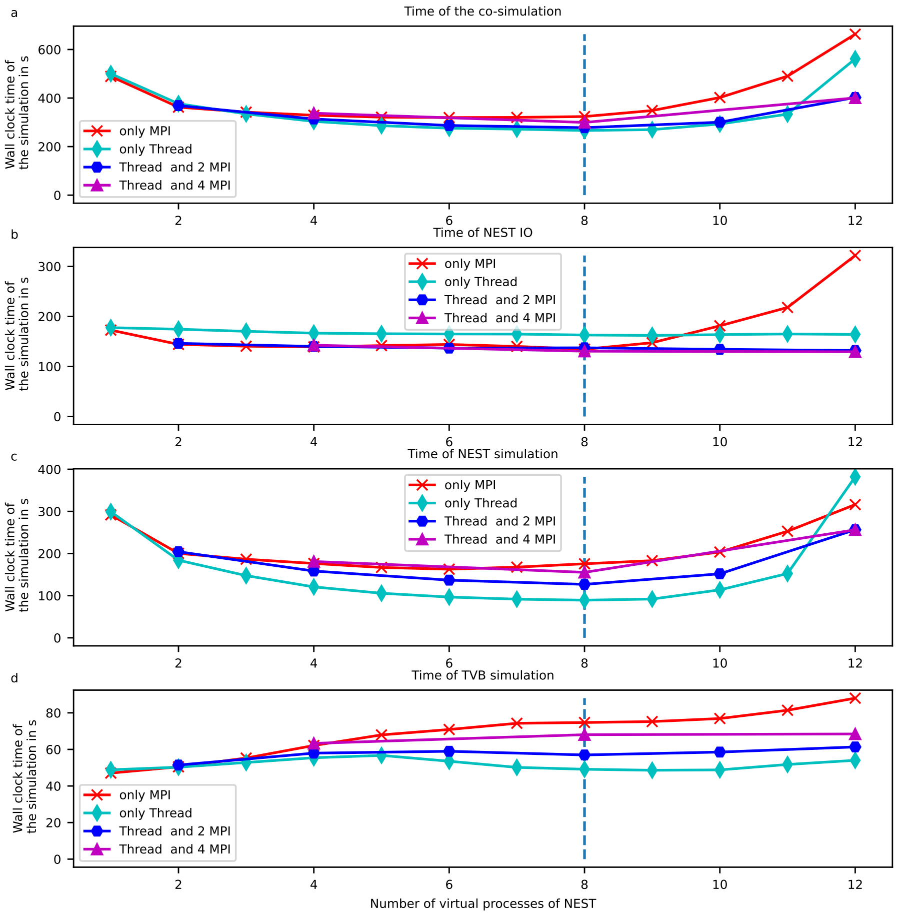


**Supplementary Figure 16** Details of the performance depending of the number of process and thread for NEST

Performance is obtained for 1 biological second on a computer (see Online Method for more details). The reference implementation use 1 MPI process, 6 virtual processes/threads, 2.0 ms to synchronize time between simulator for the simulation of 20000 neurons. Simulation time depending on the number of virtual process used by NEST. The green, blue, purple, red curves are associated with different parallelization strategy of NEST, respectively, only multithreading, 2 MPI processes with threads, 4 MPI processes with thread and only MPI processes. The horizontal green line represents the number of cores of the computer **a** The total time of the co-simulation. **b** The "IO" time of NEST **c** The "simulation" time of NEST **d** The "simulation" time of TVB

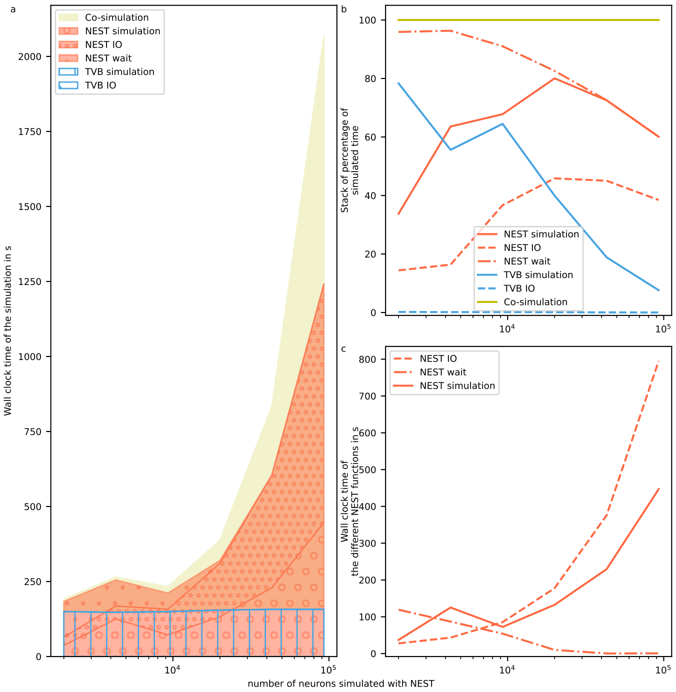


**Supplementary Figure 17** Performance of the co-simulation on one supercomputer Performance is obtained for 1 biological second on a computer. The reference implementation use 1 MPI process, 6 virtual processes/threads, 2.0 ms to synchronize time between simulator for the simulation of 20000 neurons. The node of Jusuf, the supercomputer, content 2 AMD EPYC 7742 @ 2.25 GHz * 64 cores * 2 threads, 256 (16× 16) GB DDR4 with 3200 MHz, connected by InfiniBand HDR100 (Connect-X6). The transfer modules and TVB has on one node and NEST on one or multiple other nodes. Simulation time depending on the number of neurons simulated with NEST. **a** The wall clock time of the simulator depending on the number of neurons. The total time of the co-simulation is represented in yellow. The ”simulation”, ”IO” and ”wait” times of NEST are represented in the red surface with respectively hatches with big circles, small circles and points. The ”simulation” and ”IO” times of TVB are represented in the blue surface with respectively hatches horizontal lines and oblique lines. The ”simulation” time for TVB is constant. The sum of ”simulation” and ”IO” time of NEST is higher than the TVB ”simulation”. **b** The wall clock time for the co-simulation (yellow curve), NEST (red curves) and TVB (blue curves) by the total wall clock time. The solid, dashed and dashed dotted curves are associated with ”simulation”, ”IO” and ”wait” time of NEST. The solid and dashed line is associate to ”simulation” and ”IO” time of TVB. The initialisation and configuration time increase with the number of neurons. **c** The contribution of NEST module to the total amount of the wall clock time normalizes between 0 and 100. Each contribution is reported as red curve and for increasing numbers of neurons. The solid, dashed and dashed dotted curves represent ”simulation”, ”IO” and ”wait” time of NEST respectively. The ”IO” time of NEST increases exponentially with the number of neurons and is higher than the ”simulation time when the number of neurons is higher than 6*1e4 of neurons.

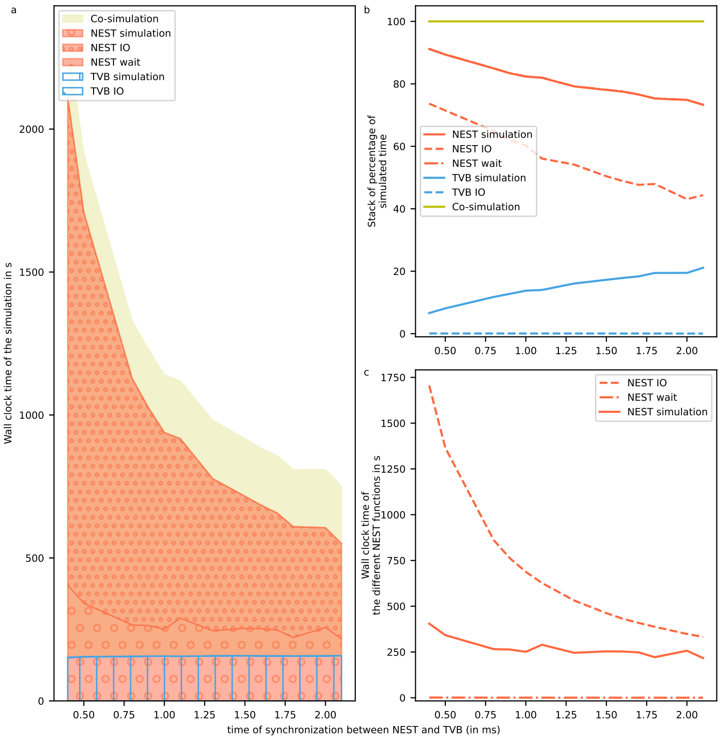


**Supplementary Figure 18** Performance of the co-simulation on one supercomputer Performance is obtained for 1 biological second on a computer. The reference implementation use 1 MPI process, 6 virtual processes/threads, 2.0 ms to synchronize time between simulator for the simulation of 20000 neurons. The node of Jusuf, the supercomputer, content 2 AMD EPYC 7742 @ 2.25 GHz * 64 cores * 2 threads, 256 (16× 16) GB DDR4 with 3200 MHz, connected by InfiniBand HDR100 (Connect-X6). The transfer modules and TVB has on one node and NEST on one or multiple other nodes. Simulation time depending on the synchronized time between simulator. **a** The wall clock time of the simulator. The simulation time reduces with the increase of the synchronization time between simulators. This reduction is due to the reduction of NEST "IO" time. The total time of the co-simulation is represented in yellow. The "simulation", "IO" and "wait" times of NEST are represented in the red surface with respectively hatches with big circles, small circles and points. The "simulation" and "IO" times of TVB are represented in the blue surface with respectively hatches horizontal lines and oblique lines. The "simulation" time for TVB is constant. The sum of "simulation" and "IO" time of NEST is higher than the TVB "simulation". **b** The wall clock time for different co-simulation modules normalized by the total wall clock time. All the curves are shown for an increase in the synchronized time between simulators. **c** The contribution of NEST module to the total amount of the wall clock time normalizes between 0 and 100. Each contribution is reported as red curve and for increasing synchronized time. The reduction follows a logarithm function.

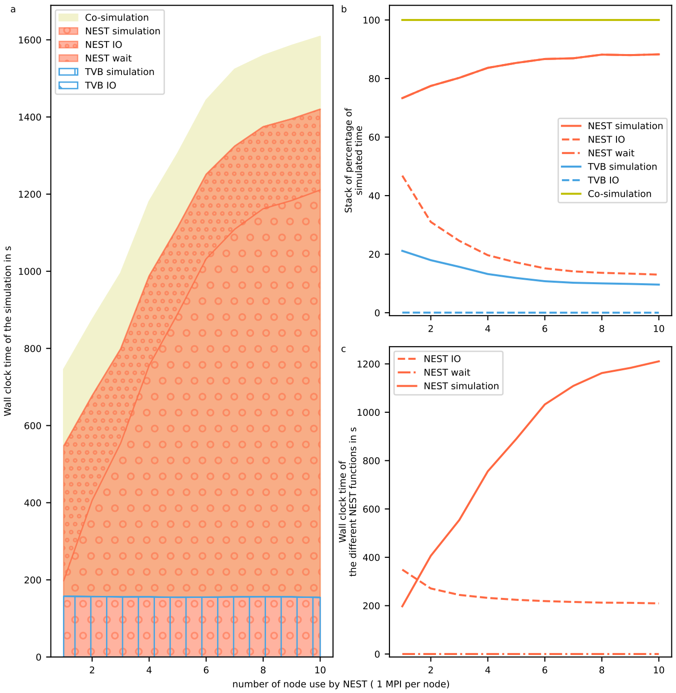


**Supplementary Figure 19** Performance of the co-simulation on one supercomputer Performance is obtained for 1 biological second on a computer. The reference implementation use 1 MPI process, 6 virtual processes/threads, 2.0 ms to synchronize time between simulator for the simulation of 20000 neurons. The node of Jusuf, the supercomputer, content 2 AMD EPYC 7742 @ 2.25 GHz * 64 cores * 2 threads, 256 (16× 16) GB DDR4 with 3200 MHz, connected by InfiniBand HDR100 (Connect-X6). The transfer modules and TVB has on one node and NEST on one or multiple other nodes. Simulation depending on the number of nodes used by NEST. **a** The wall clock time of the simulator as a function of the number of nodes used by NEST. The increase of the nodes create overhead communication in side NEST because the network is small. Moreover, the minimum delay in the network is the same as the integration step which creates an overhead of communication in NEST simulation. The wall clock time of the simulator depending on the number of neurons. The total time of the co-simulation is represented in yellow. The "simulation", "IO" and "wait" times of NEST are represented in the red surface with respectively hatches with big circles, small circles and points. The "simulation" and "IO" times of TVB are represented in the blue surface with respectively hatches horizontal lines and oblique lines. The "simulation" time for TVB is constant. The sum of "simulation" and "IO" time of NEST is higher than the TVB "simulation". **b** The wall clock time for different co-simulation modules normalized by the total wall clock time. **c** The contribution of NEST module to the total amount of the wall clock time normalized between 0 and 100. The NEST "IO" time remains constant with the increase of the number of nodes.

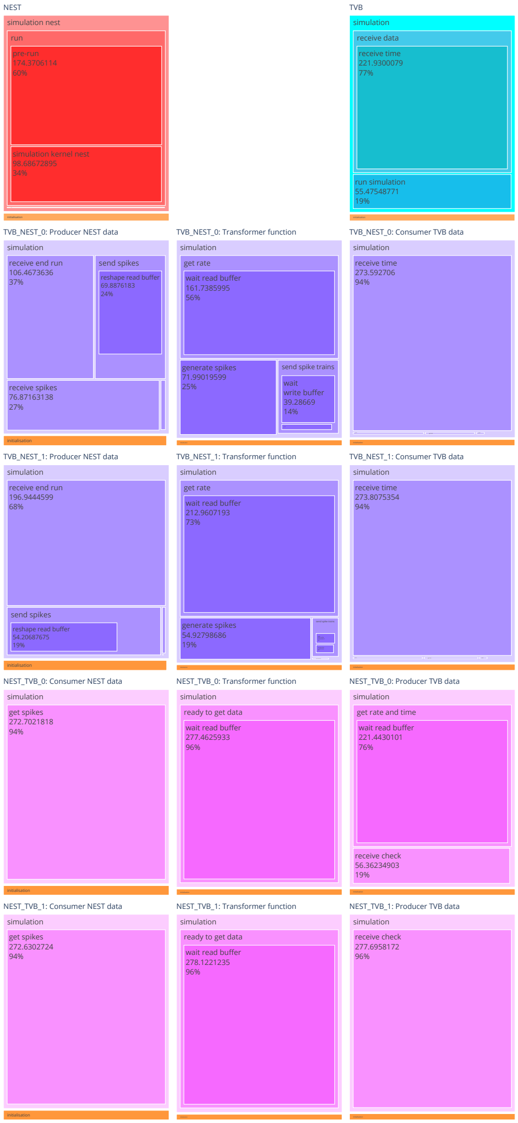


**Supplementary Figure 20** Details of the timer of one run for the reference configuration This tree-map represents the timer for each component of the transfer module and for the modules NEST and TVB at the top. The orange bar under each box represents the time required for the initialisation. Each rectangle represents the time spent on each specific piece of code.

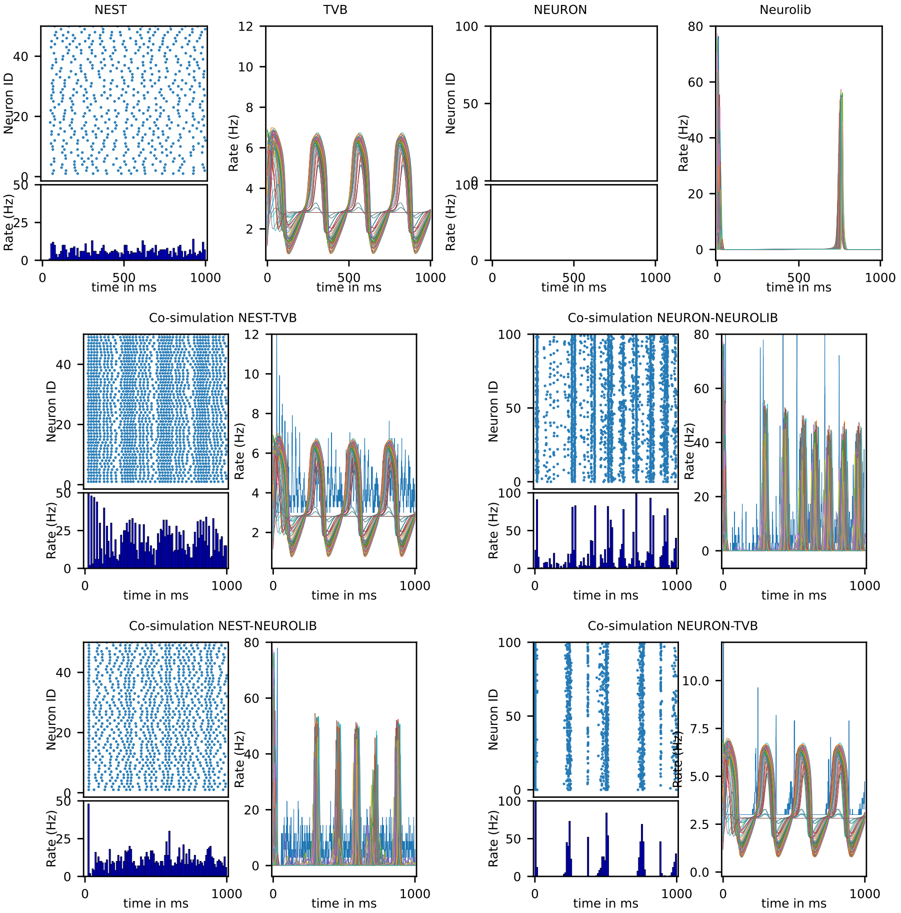


**Supplementary Figure 21** Proof of concept of replacing NEST and TVB by other simulators

For spiking neuron simulators (NEST and NEURON), the result of the simulation is, at the top, spike trains of the neurons and, at the bottom, the instantaneous firing rate of the network with a bin of 10ms. For neural mass simulators (TVB and Neurolib), the result is the firing rate of the excitatory neurons.

The first row is the results of different examples without co-simulation on different simulators. The second and third row is the result of the coupling example together simulated by using co-simulation. The co-simulation results shows the interaction of examples between them and the possibility to simulate this four different multiscale examples. The code is available here: https://github.com/multiscale-cosim/TVB-NEST-demo/tree/proof-concept

| A1 | Co-simulator environment | |
|---|---|---|
| Simulator | NEST[1] | TVB[2] |
| Version | 3.0 | 2.0 |
| Integrator method | 4th order Runge-Kutta-Fehlberg method | Heun method |
| Integration step size | 0.1 ms | 0.1 ms |
| Synchronization time step | 2.0 ms | |
| Simulated time | 60.0 s | |
| Analyzed time | between 42.5 s and 53.5 | |
| Type of I/O interface | proxy input | proxy region |

| A2 | Co-simulator architecture | |
|---|---|---|
| Reference model | The mouse brain with 104 regions | |
| Simulator | NEST | TVB |
| Number of simulated region | 2 | 102 |
| Number of MPI processes | 3 | 1 |
| Number of thread per process | 6 | 1 |
| Number of random seeds | 1 | |
| Transfer module | NEST to TVB | TVB to NEST |
| number of transfer module | 2 | 2 |
| Number of MPI processes | 2 | 2 |
| Number of threads or processes | 6 | 6 |
| Number of random seeds | 1 | 1 |

| B1 | NEST : Model Summary |
|---|---|
| Topology | left and right CA1 connected to TVB |
| Population | 2 by regions : excitatory and inhibitory |
| Connectivity | random convergent connection |
| Neuron Model | adaptive exponential leaky integrate and fire neurons[3], fixed threshold and fixed absolute refractory time |
| Synapse Model | conductance-based exponential shape |
| Plasticity | -- |
| Input | Independent fixed rate Poisson generator spike trains to all neurons and spike trains from TVB |
| Measurement | Voltage, Adaptation Current, Spike Activity and Model of Local Field Potential signal |

| B2 | NEST : Topology |
|---|---|
| regions | 2 regions ( left CA1 and right CA1) |
| number of neurons by regions | N=10000 |
| percentage of inhibitory neurons | $g_{inh}$=20% |

| B3 | NEST : Population by regions | |
|---|---|---|
| **Name** | **Elements** | **Size** |
| E | aeif_cond_exp | N_e = (1-$g_{inh}$)N = 8000 |
| I | aeif_cond_exp | N_i = $g_{inh}$N = 2000 |
| $P_{ext}$ | poisson generator | 1 |
| $I_{ext}$ | spike generator (input from TVB) | N |

| B4 | NEST : Neuron Model |
|---|---|
| Name | aeif |
| Type | adaptive exponential leaky integrator[3] and fire with conductance synapse |
| subtreshold dynamics | $$\begin{aligned} C_m \frac{dV_m}{dt} =& - g_L(V_m - E_L) + g_L \Delta_T e^{\frac{V_m - V_{th}}{\Delta_T}} \\ & - g_e(t)(V_m - E_{ex}) - g_i(t)(V_m - E_{in}) \\ & - W + I_e \\ \tau_w \frac{dW}{dt} =& a(V_m - E_L) - W \end{aligned}$$ |
| reset condition | For $t^{(f)} = \{t \mid V_m(t) >= V_{peak}\}$<br>• $V_m([t^{(f)};\ t^{(f)} + t_{ref}]) = V_{reset}$<br>• $W([t^{(f)}]) = W([t^{(f)}]) + b$ |

| B5 | NEST : Synapse Model |
|---|---|
| Name | cond_exp |
| Type | post-synaptic conductance in the form of truncated exponentials |
| Coupling equation | $$g_e(t) = \sum_{t_j^{(f)}} w_j exp\_trunc(t - t_j, \tau_{ex}) \text{ with } w_j > 0.0$$ $$g_i(t) = \sum_{t_j^{(f)}} w_j exp\_trunc(t - t_j, \tau_{in}) \text{ with } w_j < 0.0$$ $$exp\_trunc(t, \tau) = e^{1-\frac{t}{\tau}} Heaviside(t)$$ |

| B6 | NEST : Excitatory Neuron Model Parameters | | | |
|---|---|---|---|---|
| | case | Asynchronous | Irregular Synchronous | Regular bursting |
| $C_m$ | Capacity of the membrane | 200.0 pF | | |
| $t_{ref}$ | Duration of refractory period | 5.0 ms | | |
| $V_{reset}$ | Reset value for $V_m$ after a spike | -64.5 mV | -64.5 mV | -47.5 mV |
| $E_L$ | Leak reversal potential | -64.5 mV | -64.5 mV | -74.0 mV |
| $g_L$ | Leak conductance | 10.0 nS | | |
| $\Delta_T$ | Slope factor | 2.0 mV | | |
| $V_{peak}$ | Spike detection threshold | 0.0 mV | | |
| $a$ | Subthreshold adaptation | 0.0 nS | | |
| $b$ | Spike-triggered adaptation | 10.0 pA | 100.0 pA | 50.0 pA |
| $\tau_w$ | Adaptation time constant | 500.0 ms | 500.0 ms | 150.0 ms |
| $V_{th}$ | Spike initiation threshold | -50.0 mV | | |
| $I_e$ | Constant external input current | 0.0 pA | | |
| $E_{ex}$ | Excitatory reversal potential | 0.0 mV | | |
| $E_{in}$ | Inhibitory reversal potential | -80.0 mV | | |
| $V_m$ | Initialization of the voltage membrane | -64.5 mV | -64.5 mV | -47.5 mV |
| $W$ | Initialization of adaptation current | 0.0 pA | | |

| B7 | NEST : Inhibitory Neuron Model Parameters | | | |
|---|---|---|---|---|
| | case | Asynchronous | Irregular Synchronous | Regular bursting |
| $C_m$ | Capacity of the membrane | 200.0 pF | | |
| $t_{ref}$ | Duration of refractory period | 5.0 ms | | |
| $V_{reset}$ | Reset value for $V_m$ after a spike | -65.0 mV | -65.0 mV | -75.0 mV |
| $E_L$ | Leak reversal potential | -65.0 mV | -65.0 mV | -75.0 mV |
| $g_L$ | Leak conductance | 10.0 nS | | |
| $\Delta_T$ | Slope factor | 0.5 ms | | |
| $V_{peak}$ | Spike detection threshold | 0.0 mV | | |
| $a$ | Subthreshold adaptation | 0.0 nS | | |
| $b$ | Spike-triggered adaptation | 0.0 pA | | |
| $\tau_w$ | Adaptation time constant | 1.0 ms | | |
| $V_{th}$ | Spike initiation threshold | -50.0 mV | | |
| $I_e$ | Constant external input current | 0.0 pA | | |
| $E_{ex}$ | Excitatory reversal potential | 0.0 mV | | |
| $E_{in}$ | Inhibitory reversal potential | -80.0 mV | | |
| $V_m$ | Initialization of the voltage membrane | -65.0 mV | -65.0 mV | -75.0 mV |
| $W$ | Initialization of adaptation current | 0.0 pA | | |

| B8 | NEST : Connectivity between regions | | | |
|---|---|---|---|---|
| parameter synapses | | | | |
| $\tau_{ex}$ | Rise time of excitatory synaptic conductance | | | 5.0ms |
| $\tau_{in}$ | Rise time of inhibitory synaptic conductance | | | 5.0ms |
| **Name** | **Source** | **Target** | **Weights** | **Pattern** |
| EE_global | E | E I | 1.0 | Fixed total number of connection from one to another region. The number of synapses to another region is A: 1150000, IS: 3000000 and RB: 800000. The delay (161.6 ms) is defined by the multiplication of velocity (3.0 mm/ms) and distance between regions ( 53.855 mm). See for more details are on the section TVB: connectivity because delays and the weights are extract form the connectivity of TVB. |

| B9 | NEST : Connectivity inside the regions | | | |
|---|---|---|---|---|
| **Name** | **Source** | **Target** | **Weights** | **Pattern** |
| EE | E | E | 1.0 | Fixed number of input synapses ($N_e * p_{connect}$ : A and SI $400 = 8000 * 0.05$ and RB $40 = 8000 * 0.005$). The neurons can connected to itself and it can have multiple connections with the same neurons. |
| EI | E | I | 1.0 | Fixed number of input synapses ($N_e * p_{connect}$ : A and SI $400 = 8000 * 0.05$ and RB $40 = 8000 * 0.005$). The neurons can have multiple connections with the same neurons. |
| IE | I | E | g | Fixed number of input synapses ($N_i * p_{connect}$ : A and SI $100 = 2000 * 0.05$ and RB $10 = 2000 * 0.005$). The neurons can have multiple connections with the same neurons. The weight equals 10.0 for A, 5.0 for SI and 10.0 for RB. |
| II | I | I | g | Fixed number of input synapses ($N_i * p_{connect}$ : A and SI $100 = 2000 * 0.05$ and RB $10 = 2000 * 0.005$). The neurons can connected to itself and it can have multiple connections with the same neurons. The weight equals 10.0 for A, 5.0 for SI and 10.0 for RB. |

| B10 | NEST : Input | | |
|---|---|---|---|
| Poisson generator | | | |
| equation | $p(n) = \frac{\lambda^n}{n!} \exp(-\lambda)$ | | |
| implementation algorithm | Ahrens and Dieter 1982 | | |
| case | A | IS | RB |
| excitatory firing rate $\lambda_{ex}$ | 1.0 | 0.0 | 0.0 |
| inhibitory firing rate $\lambda_{in}$ | 0.0 | 0.0 | 0.0 |
| weight connection | 1.0 | 1.0 | 1.0 |
| Spike generator | | | |
| Proxy for the input of the region simulated with TVB. (see the section transformation TVB to NEST) | | | |

| B11 | NEST : Measurement (part 1) | | |
|---|---|---|---|
| state variable | Voltage membrane, adaptation current | precision | 0.1 |
| | | number of recorded neurons | 10 excitatory and 10 inhibitory |
| | spike time | precision | 0.1 ms |
| | | number of recorded neurons | all |
| spike activities | raster plot | precision | 0.1 ms |
| | histogram of instantaneous firing rate | bins | 0.1ms |
| | simple moving average | windows size | T (20ms) |
| | spectogram | method | Welch's method |
| | | sampling frequencies | $10^4$Hz |
| | | window shape | Hann window |
| | | length of each segment | $10^4$ |
| | | length of the FFT | $10^4$ |
| | | number of points overlapping | $5.10^3$ |
| | | detrend | removing the mean |
| | | sides | only real part |

| B11 | NEST : Measurement (part 2) | |
|---|---|---|
| Micro -electodes: Local Field Potential | software<br>number of MPI<br>number random seed | HybridLFPy[4]<br>2<br>2 |
| | number of segment by neuron | defined by the method lambda100 of Neuron |
| | resolution | 0.1 ms |
| | soma position | random in a cylinder of radius 2000 mm and height of 100mm with a minimal distance of 1mm. The center of the cylinder is (0,-400). |
| | excitatory neurons | |
| | morphology | pyramidal cell of Shuman 2020 [5] without biophysics and synapses mechanisms |
| | initial membrane potential | $V_m$ (-64.5 mV or -47.5 mV) |
| | axial resistance | 150.0 Ohm |
| | membrane capacitance | $C_m$ ( 200pF ) |
| | passive mechanism | yes |
| | passive reversal potential | $E_L$ (-64.5 mV or -74.0 mV)) |
| | passive conductance | $g_L$ (10 nS) |
| | inhibitory neurons | |
| | morphology | basket cell of Shuman 2020 [5] without biophysics and synapses mechanisms |
| | initial membrane potential | $V_m$ (-64.5 mV or -75.0 mV) |
| | axial resistance | 150.0 Ohm |
| | membrane capacitance | $C_m$ ( 200pF ) |
| | passive mechanism | yes |
| | passive reversal potential | $E_L$ (-65.0 mV or -75.0 mV)) |
| | passive conductance | $g_L$ (10 nS) |

<table>
<tr><th>B11</th><th colspan="2">NEST : Measurement(part 3)</th></tr>
<tr><td rowspan="17">Micro<br>-electodes:<br>Local Field<br>Potential</td><td colspan="2">connectivity</td></tr>
<tr><td>layers</td><td>2 : [[300,-100],[-100,-600]]</td></tr>
<tr><td>synapse shape</td><td>truncated exponential</td></tr>
<tr><td>delay and weight distribution</td><td>homogeneous values by population</td></tr>
<tr><td>excitatory connection by layers and populations</td><td><table><tr><td>$N_e * p_{connect} * 0.5$</td><td>0</td></tr><tr><td>$N_e * p_{connect} * 0.5$</td><td>$N_i * p_{connect}$</td></tr></table></td></tr>
<tr><td>excitatory weight</td><td>1.0</td></tr>
<tr><td>excitatory delay</td><td>dt (0.1 ms)</td></tr>
<tr><td>inhibitory connection by layers and populations</td><td><table><tr><td>0</td><td>0</td></tr><tr><td>$N_e * p_{connect}$</td><td>$N_i * p_{connect}$</td></tr></table></td></tr>
<tr><td>inhibitory weight</td><td>g ( 5.0 or 10.0)</td></tr>
<tr><td>inhibitory delay</td><td>dt (0.1 ms)</td></tr>
<tr><td colspan="2">electrodes</td></tr>
<tr><td>extracellular conductivity</td><td>0.3 S</td></tr>
<tr><td>electrode positions and contacts surface normal</td><td><table>
<tr><th colspan="3">positions</th><th colspan="3">normal</th></tr>
<tr><th>x</th><th>y</th><th>z</th><th>x</th><th>y</th><th>z</th></tr>
<tr><td>1273</td><td>1273</td><td>1273</td><td>1</td><td>1</td><td>0</td></tr>
<tr><td>1273</td><td>-1273</td><td>-1273</td><td>1</td><td>1</td><td>0</td></tr>
<tr><td>-1273</td><td>-1273</td><td>15</td><td>1</td><td>1</td><td>0</td></tr>
<tr><td>-15</td><td>15</td><td>-15</td><td>1</td><td>1</td><td>0</td></tr>
<tr><td>1288</td><td>1258</td><td>1288</td><td>1</td><td>-1</td><td>0</td></tr>
<tr><td>1258</td><td>1288</td><td>1258</td><td>1</td><td>-1</td><td>0</td></tr>
<tr><td>1288</td><td>1258</td><td>-1800</td><td>1</td><td>-1</td><td>0</td></tr>
<tr><td>-1800</td><td>-1800</td><td>-1800</td><td>1</td><td>-1</td><td>0</td></tr>
<tr><td>-385</td><td>-385</td><td>-415</td><td>1</td><td>0</td><td>0</td></tr>
<tr><td>-415</td><td>-385</td><td>-385</td><td>1</td><td>0</td><td>0</td></tr>
<tr><td>-415</td><td>-415</td><td>-385</td><td>1</td><td>0</td><td>0</td></tr>
<tr><td>-385</td><td>-415</td><td>-415</td><td>1</td><td>0</td><td>0</td></tr>
</table></td></tr>
<tr><td>contact shape</td><td>circle of radius 20 mm</td></tr>
<tr><td>number of discrete point for compute the average potential</td><td>20</td></tr>
<tr><td>assumption method</td><td>soma as point</td></tr>
</table>

| C1 | TVB : Model Summary |
|---|---|
| Neural Mass model | Mean Adaptive Exponential |
| Connectivity | Mouse connectome with 104 regions |
| Coupling | linear coupling |
| stimulus | -- |
| Monitors | ECoG (Electrocorticography) and state variable of the mean field |

| C2 | TVB : Coupling |
|---|---|
| Name | Linear |
| Type | Linear coupling |
| equations | $\nu_{ext_k} = a * \left( \sum_{104}^{j=1} u_{kj} \nu_{e_j}(t - \tau_{kj}) \right) + b$ where $u_{kj}$ are the elements of the weights matrix, $\tau_{kj}$ are the elements of the delay matrix and $\nu_{e_j}$ are the mean excitatory firing rate of the regions j. |
| parameters | $a = 1.0$ and $b = 0.0$ |

| C3 | TVB : Connectivity |
|---|---|
| Connectivity is extracted from tracer data as explained by the paper TVBM[6] | |
| number of region | 104 |
| tract lengths | maximum : 115.46 and mean : 53.58 |
| speed | 3 ms |
| weights | The weigths are normalize such as the sum of the input weight to one region eguals 1. (maximum : 0.73 and mean : 0.02) |
| centers | average center of mouse brain : [57., 74.97, 42.53] |
| orientation | the orientation is defined by a vector from the average center of mouse brain to the center of the regions |

| C3 | TVB : Connectivity |
|---|---|
| region name | the region name are extracted from Allen Mouse Brain Connectivity Atlas (17/01/2017)[7] |
| | Right Primary motor area, Right Secondary motor area, Right Primary somatosensory area nose, Right Primary somatosensory area barrel field, Right Primary somatosensory area lower limb, Right Primary somatosensory area mouth, Right Primary somatosensory area upper limb, Right Supplemental somatosensory area, Right Gustatory areas, Right Visceral area, Right Dorsal auditory area, Right Primary auditory area, Right Ventral auditory area, Right Primary visual area, Right Anterior cingulate area dorsal part, Right Anterior cingulate area ventral part, Right Agranular insular area dorsal part, Right Retrosplenial area dorsal part, Right Retrosplenial area ventral part, Right Temporal association areas, Right Perirhinal area, Right Ectorhinal area, Right Main olfactory bulb, Right Anterior olfactory nucleus, Right Piriform area, Right Cortical amygdalar area posterior part, Right Field CA1, Right Field CA3, Right Dentate gyrus, Right Entorhinal area lateral part, Right Entorhinal area medial part dorsal zone, Right Subiculum, Right Caudoputamen*, Right Nucleus accumbens*, Right Olfactory tubercle*, Right Substantia innominata*, Right Lateral hypothalamic area*, Right Superior colliculus sensory related*, Right Inferior colliculus*, Right Midbrain reticular nucleus*, Right Superior colliculus motor related*, Right Periaqueductal gray*, Right Pontine reticular nucleus caudal part*, Right Pontine reticular nucleus*, Right Intermediate reticular nucleus*, Right Central lobule*, Right Culmen*, Right Simple lobule*, Right Ansiform lobule*, Right Paramedian lobule*, Right Copula pyramidis*, Right Paraflocculus*, Left Primary motor area, Left Secondary motor area, Left Primary somatosensory area nose, Left Primary somatosensory area barrel field, Left Primary somatosensory area lower limb, Left Primary somatosensory area mouth, Left Primary somatosensory area upper limb, Left Supplemental somatosensory area, Left Gustatory areas, Left Visceral area, Left Dorsal auditory area, Left Primary auditory area, Left Ventral auditory area, Left Primary visual area, Left Anterior cingulate area dorsal part, Left Anterior cingulate area ventral part, Left Agranular insular area dorsal part, Left Retrosplenial area dorsal part, Left Retrosplenial area ventral part, Left Temporal association areas, Left Perirhinal area, Left Ectorhinal area, Left Main olfactory bulb, Left Anterior olfactory nucleus, Left Piriform area, Left Cortical amygdalar area posterior part, Left Field CA1, Left Field CA3, Left Dentate gyrus, Left Entorhinal area lateral part, Left Entorhinal area medial part dorsal zone, Left Subiculum, Left Caudoputamen*, Left Nucleus accumbens*, Left Olfactory tubercle*, Left Substantia innominata*, Left Lateral hypothalamic area*, Left Superior colliculus sensory related*, Left Inferior colliculus*, Left Midbrain reticular nucleus*, Left Superior colliculus motor related*, Left Periaqueductal gray*, Left Pontine reticular nucleus caudal part*, Left Pontine reticular nucleus*, Left Intermediate reticular nucleus*, Left Central lobule*, Left Culmen*, Left Simple lobule*, Left Ansiform lobule*, Left Paramedian lobule*, Left Copula pyramidis*, Left Paraflocculus* |
| cortical region | all the region name ending by a '*' are not cortical regions |

| C4 | TVB : Neural Mass Model (part 1) |
|---|---|
| Name | Mean Ad Ex[8] |
| Type | Neural mass model of network of adaptive exponential integrate and fire excitatory and inhibitory neurons of second statistical order with adaptation |
| equation | $$\begin{aligned} T\,\frac{\partial \nu_e}{\partial t} =& (\mathcal{F}_e - \nu_e) + \frac{1}{2}\,c_{ee}\,\frac{\partial^2 \mathcal{F}_e}{\partial \nu_e \partial \nu_e} \\ & + \frac{1}{2}\,c_{ei}\,\frac{\partial^2 \mathcal{F}_e}{\partial \nu_e \partial \nu_i} + \frac{1}{2}\,c_{ie}\,\frac{\partial^2 \mathcal{F}_e}{\partial \nu_i \partial \nu_e} + \frac{1}{2}\,c_{ii}\,\frac{\partial^2 \mathcal{F}_e}{\partial \nu_i \partial \nu_i} \\ T\,\frac{\partial \nu_i}{\partial t} =& (\mathcal{F}_i - \nu_i) + \frac{1}{2}\,c_{ee}\,\frac{\partial^2 \mathcal{F}_i}{\partial \nu_e \partial \nu_e} \\ & + \frac{1}{2}\,c_{ei}\,\frac{\partial^2 \mathcal{F}_i}{\partial \nu_e \partial \nu_i} + \frac{1}{2}\,c_{ie}\,\frac{\partial^2 \mathcal{F}_i}{\partial \nu_i \partial \nu_e} + \frac{1}{2}\,c_{ii}\,\frac{\partial^2 \mathcal{F}_i}{\partial \nu_i \partial \nu_i} \\ T\,\frac{\partial c_{ee}}{\partial t} =& (\mathcal{F}_e - \nu_e)\,(\mathcal{F}_e - \nu_e) + c_{ee}\frac{\partial \mathcal{F}_e}{\partial \nu_e} + c_{ee}\frac{\partial \mathcal{F}_e}{\partial \nu_e} + c_{ei}\frac{\partial \mathcal{F}_i}{\partial \nu_e} \\ & + c_{ie}\frac{\partial \mathcal{F}_i}{\partial \nu_e} - 2c_{ee} + \frac{\mathcal{F}_e\,(1/T - \mathcal{F}_e)}{N_e} \\ T\,\frac{\partial c_{ei}}{\partial t} =& (\mathcal{F}_e - \nu_e)\,(\mathcal{F}_i - \nu_i) + c_{ee}\frac{\partial \mathcal{F}_e}{\partial \nu_e} \\ & + c_{ei}\frac{\partial \mathcal{F}_e}{\partial \nu_i} + c_{ei}\frac{\partial \mathcal{F}_i}{\partial \nu_e} + c_{ii}\frac{\partial \mathcal{F}_i}{\partial \nu_i} - 2c_{ei} \\ T\,\frac{\partial c_{ie}}{\partial t} =& (\mathcal{F}_i - \nu_i)\,(\mathcal{F}_e - \nu_e) + c_{ie}\frac{\partial \mathcal{F}_e}{\partial \nu_i} \\ & + c_{ee}\frac{\partial \mathcal{F}_e}{\partial \nu_e} + c_{ii}\frac{\partial \mathcal{F}_i}{\partial \nu_i} + c_{ie}\frac{\partial \mathcal{F}_i}{\partial \nu_e} - 2c_{ie} \\ T\,\frac{\partial c_{ii}}{\partial t} =& (\mathcal{F}_i - \nu_i)\,(\mathcal{F}_i - \nu_i) + c_{ie}\frac{\partial \mathcal{F}_e}{\partial \nu_i} + c_{ei}\frac{\partial \mathcal{F}_e}{\partial \nu_i} + c_{ii}\frac{\partial \mathcal{F}_i}{\partial \nu_i} \\ & + c_{ii}\frac{\partial \mathcal{F}_i}{\partial \nu_i} - 2c_{ii} + \frac{\mathcal{F}_i\,(1/T - \mathcal{F}_i)}{N_i} \\ \tau_{W_e}\frac{\partial W_e}{\partial t} &= -W_e + b_e\nu_e + a_e(\mu_V(\nu_e, \nu_i, \nu_{ext}, W_e) - EL_e) \\ \tau_{W_i}\frac{\partial W_i}{\partial t} &= -W_i + b_i\nu_i + a_i(\mu_V(\nu_e, \nu_i, \nu_{ext}, W_i) - EL_i) \end{aligned}$$ |
| noise equation | Ornstein-Uhlenbeck process : $\tau_{ou}\frac{dou_t}{dt} = (\mu - ou_t) + \sigma\, dW_t$ with $W_t$ is a Wiener process |

| C4 | TVB : Neural Mass Model (part 2) |
|---|---|
| transfer function | $$\begin{aligned} \mathcal{F}_e =& \mathcal{F}((\nu_e + 1e-6) + w_\sigma ou_t, \nu_{ext}, \nu_i, W_e) \\ \mathcal{F}_i =& \mathcal{F}((\nu_e + 1e-6) + w_\sigma ou_t, \nu_{ext}, \nu_i, W_i) \\ \mathcal{F} =& \frac{1}{2\,\tau_V} \cdot Erfc(\frac{V_{thre}^{eff} - \mu_V}{\sqrt{2}\,\sigma_V}) \\ V_{thre}^{eff}(\mu_V, \sigma_V, \tau_V^N =& \tau_V \frac{g_L}{Cm}) = P_0' + \sum_{x \in \{\mu_V, \sigma_V, \tau_V^N\}} P_x \cdot \left(\frac{x - x^0}{\delta x^0}\right) \\ &+ \sum_{x,y \in \{\mu_V, \sigma_V, \tau_V^N\}^2} P_{xy} \cdot \left(\frac{x - x^0}{\delta x^0}\right) \left(\frac{y - y^0}{\delta y^0}\right) \\ \mu_G(\nu_e, \nu_{ext}, \nu_i) =& ((\nu_e K_e + \nu_{ext} K_{ext}) \tau_e Q_e) + (\nu_i K_i \tau_i Q_i) + g_L \\ \mu_{V_s}(\nu_e, \nu_{ext}, \nu_i, w, \mu_G) =& \frac{((\nu_e K_e + \nu_{ext} K_{ext}) \tau_e Q_e) E_e}{\mu_G} \\ &+ \frac{(\nu_i K_i \tau_i Q_i) E_i + g_L EL_s - w}{\mu_G} \\ \sigma_V(\mu_V, \mu_G) =& \sqrt{\sum_{s \in \{e,i\}} K_s \nu_s \frac{\left(\frac{Q_s}{\mu_G}(E_s - \mu_V)\tau_s\right)^2}{2\frac{C_m}{\mu_G} + \tau_s}} \\ \tau_V(\mu_V, \mu_G) =& \frac{\sum_{s \in \{e,i\}} K_s \nu_s \left(\frac{Q_s}{\mu_G}(E_s - \mu_V)\tau_s\right)^2}{\sum_{s \in \{ex,in\}} K_s \nu_s \frac{\left(\frac{Q_s}{\mu_G}(E_s - \mu_V)\tau_s\right)^2}{2\frac{Cm}{\mu_G} + \tau_s}} \end{aligned}$$ |

| C5 | TVB : Neural Mass Model Parameters(part 1) | | | |
|---|---|---|---|---|
| | case | Asynchronous | Irregular Synchronize | Regular bursting |
| $T$ | time resolution of the mean field | 20.0ms | | |
| $C_m$ | Capacity of the membrane | 200.0 pF | | |
| $EL_e$ | Leak reversal potential excitatory($E_L$) | -64.5 mV | -64.5 mV | -74.0 mV |
| $EL_i$ | Leak reversal potential inhibitory($E_L$) | -65.0 mV | -65.0 mV | -75.0 mV |
| $g_L$ | Leak conductance | 10.0 nS | | |
| $a_e$ | Subthreshold adaptation of excitatory neurons($a$) | 0.0 nS | | |
| $b_e$ | Spike-triggered adaptation of excitatory neurons($b$) | 10.0 pA | 100.0 pA | 50.0 pA |
| $\tau_{W_e}$ | Adaptation time constant of excitatory neurons($\tau_w$) | 500.0 ms | 500.0 ms | 150.0 ms |
| $a_i$ | Subthreshold adaptation of inhibitory neurons($a$) | 0.0 nS | | |
| $b_i$ | Spike-triggered adaptation inhibitory neurons($b$) | 0.0 pA | | |
| $\tau_{W_i}$ | Adaptation time constant of inhibitory neurons($\tau_w$) | 1.0 ms | | |
| $E_e$ | Excitatory reversal potential($E_{ex}$) | 0.0 mV | | |
| $\tau_e$ | Rise time of excitatory synaptic conductance($\tau_{ex}$) | 5.0 ms | | |
| $Q_e$ | excitatory quantal conductance | 1.0 nS | | |
| $E_i$ | Inhibitory reversal potential($E_{in}$) | -80.0 mV | | |
| $\tau_i$ | Rise time of inhibitory synaptic conductance($\tau_{in}$) | 5.0 ms | | |
| $Q_i$ | inhibitory quantal conductance | 10.0 nS | 5.0 nS | 10.0 nS |
| $p_{connect}$ | probability of connection | 0.05 | 0.05 | 0.005 |
| $N_{tot}$ | Number of total neurons | 10000 | | |
| $p_i$ | percentage of inhibitory neurons | 0.2 | | |
| $N_e$ | Number of excitatory neurons | $N_{tot}(1-p_i)$ =8000 | | |
| $N_i$ | Number of inhibitory neurons | $N_{tot}p_i$ =2000 | | |

| C5 | TVB : Neural Mass Model Parameters(part 2) | | | | |
|---|---|---|---|---|---|
| $K_e$ | mean number of input excitatory synapses : $N_e p_{connect}$ | 400 | 400 | 40 | |
| $K_i$ | mean number of input inhibitory synapses : $N_i p_{connect}$ | 100 | 100 | 10 | |
| $Kext_e$ | number of external excitatory synapse | 115 | 300 | 80 | |
| $P_e$ | second order polynomial of the phenomenological threshold for inhibitory neuron in mV | $P_0$ | $P_{\mu_V}$ | $P_{\sigma_V}$ | $P_{\tau_V^N}$ |
| | | -0.0498 | 0.00506 | -0.025 | 0.0014 |
| | | $P_{\mu_V^2}$ | $P_{\sigma_V^2}$ | $P_{(\tau_V^N)^2}$ | |
| | | -0.00041 | 0.0105 | -0.036 | |
| | | $P_{\mu_V \sigma_V}$ | $P_{\mu_V \tau_V^N}$ | $P_{\sigma_V \tau_V^N}$ | |
| | | 0.0074 | -0.0012 | -0.0407 | |
| $P_i$ | second order polynomial of the phenomenological threshold for inhibitory neuron in mV | $P_0$ | $P_{\mu_V}$ | $P_{\sigma_V}$ | $P_{\tau_V^N}$ |
| | | -0.0514 | 0.004 | -0.0083 | 0.0002 |
| | | $P_{\mu_V^2}$ | $P_{\sigma_V^2}$ | $P_{(\tau_V^N)^2}$ | |
| | | -0.0005 | 0.0014 | -0.014 | |
| | | $P_{\mu_V \sigma_V}$ | $P_{\mu_V \tau_V^N}$ | $P_{\sigma_V \tau_V^N}$ | |
| | | 0.0045 | 0.0028 | -0.00153 | |
| $\nu_{ext}$ | external input | see coupling section | | | |
| $w_\sigma$ | weight of the noise | 0.0002 | 0.0006 | 0.002 | |
| $\sigma$ | variation of the noise | 0.2 | | | |
| $\mu$ | mean of the noise | 0.0 | | | |
| $\tau_{ou}$ | mean of the noise | 20.0 | | | |
| | initial condition (random between maximum and minimum) | $\mu_E(kHz) : (0., 0.)$ $\mu_i(kHz) : (0., 0.)$ $c_{ee} : (0., 0.)$ $c_{ei} : (0., 0.)$ $c_{ii} : (0., 0.)$ $W_e(pA) : (0., 5.)$ $W_i(pA) : (0., 0.)$ | | | |

<table>
<tr><th>C6</th><th colspan="2">TVB : Monitor</th></tr>
<tr><td></td><td>proxy node</td><td>node</td></tr>
<tr><td>state variable</td><td>Only the mean firing rate of the excitatory population because it's the coupling variable and it's extract from NEST simulation</td><td>mean firing rate of excitatory and inhibitory population, the variation of excitatory and inhibitory firing rate, the co-variation between excitatory and inhibitory firing rate, mean adaptive current of excitatory and inhibitory firing rate</td></tr>
<tr><td></td><td>precision</td><td>dt (0.1ms)</td></tr>
<tr><td rowspan="2">SEEG</td><td>equation</td><td>$\Psi_{ECoG}(channel, t) = P.\nu_e + noise$<br>where P is the gain matrix and N is the mean firing rate of excitatory population<br>$P_{ij} = scaling\_factor * region\_volume_j / r_i - r_j$ where $r_i$ is the position of the contact point of the channel and $r_j$ is the center of region j.</td></tr>
<tr><td>contact position</td><td>
<table>
<tr><th></th><th>x</th><th>y</th><th>z</th></tr>
<tr><td rowspan="8">left hemisphere</td><td>40.0</td><td>80.0</td><td>79.5</td></tr>
<tr><td>20.0</td><td>80.0</td><td>72.0</td></tr>
<tr><td>30.0</td><td>70.0</td><td>76.5</td></tr>
<tr><td>30.0</td><td>90.0</td><td>75.5</td></tr>
<tr><td>22.5</td><td>72.5</td><td>73.0</td></tr>
<tr><td>22.5</td><td>87.5</td><td>72.5</td></tr>
<tr><td>37.5</td><td>72.5</td><td>78.5</td></tr>
<tr><td>37.5</td><td>87.5</td><td>78.5</td></tr>
<tr><td rowspan="8">right hemisphere</td><td>94.0</td><td>80.0</td><td>69.</td></tr>
<tr><td>74.0</td><td>80.0</td><td>78.5</td></tr>
<tr><td>84.0</td><td>70.0</td><td>74.5</td></tr>
<tr><td>84.0</td><td>90.0</td><td>74.0</td></tr>
<tr><td>76.5</td><td>72.5</td><td>77.5</td></tr>
<tr><td>76.5</td><td>87.5</td><td>77.5</td></tr>
<tr><td>91.5</td><td>72.5</td><td>70.5</td></tr>
<tr><td>91.5</td><td>87.5</td><td>70.</td></tr>
</table>
</td></tr>
<tr><td></td><td>$scaling\_factor$</td><td>1.0</td></tr>
<tr><td></td><td>$region\_volume$</td><td>The volume is extract from the volume mapping of Allen Mouse Brain Connectivity Atlas<br>mean: 3712.29 max: 16245.0 min: 957.0</td></tr>
</table>

| D | Transformation NEST to TVB : model |
|---|---|
| Name | SMFR : sliding mean firing rate |
| Type | sliding mean over the histogram of the instantaneous firing rate |
| Input | spike trains of excitatory neurons from one brain region for synchronized time (2ms) |
| Output | mean firing rate of excitatory population of the brain region for synchronized time (2ms) |
| equation | $\forall t \geq T, \mathrm{SMFR}(t) = \frac{\sum_{s=t-T}^{t} \sum_{n=1}^{N_e} spike(n,s)}{N_e T} * 10^3 \text{ (KHz)}$ <br> where $spike(n,s) = \begin{cases} 1 \text{ if neuron n create spike at time s} \\ \quad \text{with a presicion of dt} \\ 0 \text{ else} \end{cases}$ |
| parameters | size of the windows $T$ : 20.0 ms ( same as TVB) |
| parameters | number of neurons $N_e$ : 8000 ( same as NEST) |
| parameters | precision of the integration $dt$ : 0.1 ms (same as TVB and NEST) |
| initialization | The transfer module doesn't have initialized because TVB used its initialization for starting the communication. |

| E | Transformation TVB to NEST : model |
|---|---|
| Name | MIP |
| Type | Multiple Interaction Process[9] |
| Input | incoming excitatory firing rate of a brain region for synchronized time (2ms) |
| Output | spike trains to individual neurons with a correlation of $p$ for synchronized time (2ms) |
| equation | reference spike train:<br>$x_{ref}(t) = InhomogenousPoissonProcess((\nu_{input}(t)nb_{synapse} + 1e - 12)/p)$<br>input individual spike train to the neuron n :<br>$x_n(t) = x_{ref}B(size(x_{ref}), p)$ |
| where $\nu_{input}$ | mean external excitatory firing rate computed by TVB for the NEST population. |
| $nb_{synapse}$ | number of external input synapse (A:115, IS:300, RB:80) |
| $p$ | percentage of shared neurons (A:0.01, IS:0.1, RB:0.01) |
| $B$ | binomial law |
| The implementation of the Inhomogenous Poisson Process | Dedicated function from the python library elephant (version 0.9), which : 1) generates spike trains with homogeneous Poisson generator for the highest rate; 2) removes some spikes for having rate variation based on the input rates. The homogeneous Poisson generator computes the time interval between each spike using the exponential random generator of numpy. |
| initialization | The initial rate send to TVB are zeros during the first $t_{synch}$ (2ms). |

### Supplementary Note 1| Guidelines Input/Output (I/O) interface

Before the creation of the I/O interface, the simulator has to be analysed. The objective is to verify the existence of output and input devices, the paradigm of parallelization, the tools for the parallelization and the properties to keep. This analysis will help to modify the architecture of the simulator. The modification needs to be in accordance with the simulator development and its maintainability. The last part is the creation of a wrapper to communicate with the transfer module if it is required. Two paragraphs will provide further details about how to transfer data from NEST to TVB and vice versa.

#### NEST I/O interface

NEST has 2 types of devices : stimulating and recording devices. These devices receive or send messages mainly based on spikes times. The parallelization use MPI and/or threading depending on its parametrization and it is based principally on events transfer (spikes between neurons). The important property to conserve is its scalability. From this analysis, a new interface is integrated in the version 3 of NEST and use MPI communication. The modification architecture of NEST is the creation of specific back end of the recording and stimulating devices and the reformatting of input devices to include the usage of specific back end. Each back-end use a communication protocol (see supplementary figures 3) which include the transmission of the state of NEST with tags and the exchange of data. This interface is directly used by the transfer module(see supplementary figures 10).

#### TVB I/O interface

The TVB simulator presents monitor classes for recording values and stimulating classes. There is not parallelization optimization and everything is storage in memory (recording simulated data, stimulating and transfer data between Neural Mass). This simulator does not have specific properties to keep for the optimization of the simulation. The prototype uses a new monitor which modifies the simulator dynamically. This new monitor adds an extra buffer to delay the simulated data. The advantage of it is the possibility to use proxy nodes and an IO interface to include external data during this delay time. To be in accordance with the simulator development and the maintainability of the simulator, this new monitor is not included in the simulator but a new class of simulator will be created for the simulation. This interface is not enough for the communication with the transfer module because there is a need to use MPI communication. A wrapper around this interface is implemented to overcome this requirement(see supplementary figure 7 for details). A bug present in the implementation monitor does not take into account the time of synchronization between the simulator but it does not have an impact on the co-simulation dynamic.

### Supplementary Note 2| Guideline for implementation of transfer module

This section focus on the intention behind the implementation of the transfer modules. In the future, the transfer module will be improved and used by two types of scientists. The neuroscientist or physician will modify it for creating new models and adapted it to their scientific questions. In parallel, computational scientists will work for improving the speed of communication between all modules and components. Furthermore, in the future, there will be a need to add other simulators as Neuron, Arbor, NRP,... The architecture design needs to simplify the addition of other simulators and other types of data (membrane voltage, current, ...).

The separation of the neuroscience research and computer science research is done by the separation of the functions of the transfer module in three components/objects/processes: two for the I/O interface with simulators and one for transformation functions (see Supplementary Figure 11). A neuroscientist will modify principally the transformation components where the meaning of the transformation of data is required and important for his work. A computational scientist will focus on the optimization of the communication with the interface with a simulator, the internal communication and the management of the flux of data.

For avoiding conflict between this type of research, there is a simple API for receiving and sending data in each component (see examples of activity of diagram of the transfer components in the Supplementary Figure 13). The only constraint to the neuroscientist is to respect the buffering of data in the transformation function by releasing the input connection before the access to the output connection. Moreover this simple API is implemented following the abstract factory pattern. This design pattern is chosen to help the comparison of different implementations of communication and the integration of new simulators.

The API address partially the constraint of the simplification for adding a new simulator because there is only missing part is component for the interface with the simulator, the rest can be reused. The other element of the architecture for the respect of this constraint is the separation of files for each simulator and the encapsulation of the interface in abstract class following a composite pattern.

The second constraint of the simplification of adding a new type of data is respected by the imposition of a convention for the management of data. This convention is composed of four functions and one Boolean for sending data and the same for receiving data. The functions are ”ready for transfer data?”, ”transfer the data”, ”end of transfer data” and ”release the connection”. The Boolean contains if the connection is closed by the other side or not.

### Supplementary Note 3| Detail characterization of the workflow TVB-NEST

This characterization is based on the taxonomies proposed in Gomes 2018.[10]. However, this taxonomy is not the best for this workflow because the transformation modules are not taking into account and one hypothesis of this taxonomy is the presence of an launcher which is not the case for the workflow.

#### Non-Functional Requirements

- Fault tolerance : No (NEST does not store the previous state and the communication spike which create the impossibility to coming back in the future)
- Configuration reused : Yes (the configuration of each simulator is independent and defines during the initialization)
- Performance : Yes and No (the scalability and the parallelization of the simulator is kept but there is not the modulation of the integration step or signal extrapolation).
- IP Protection : No protection (NEST and TVB do not use protected models.)
- Parallelism : Yes (the communication use MPI and each simulator is run in individual processes)
- Distributed : Yes (the workflow keep the properties of NEST to be simulated in distributed way.)
- Hierarchy : Yes (the workflow is independent of the model for each simulator and the transformation function. There is some requirement for the connection between modules which create the dependencies.)
- Scalability : No (it is dependent on the simulators)
- Platform independent : Yes and No (it requires some dependence on the platform but it can be passed by the usage of docker or singularity)
- Extensibility : Yes and No (there are some extra modules as NESTML or TVB which can create models for each simulator but not a specific extension for the transformation and all the simulations.)
- Accuracy : No (there are any simulators which provide the errors or the convergence of the simulations.)
- Open Source : Yes (each simulator is open source and the workflow is also open source)

#### Simulator Requirements

##### Information Exposed

- Frequency of State : No (the frequency of the state for the simulator and the co-simulation is fixed during the initialization)
- Frequency of Outputs: No (same as before. Moreover, TVB can have an output frequency lower than this internal integration frequency)
- Detailed Model : Yes (the code for all the models is available)
- Nominal Values of Outputs : dependent of the output and the models used

- Nominal Values of State : dependent on the model
- I/O Signal Kind : No (there are not a master algorithm but NEST has some internal statement about the signal communicate between devices and nodes.)
- Time Derivative : Output only
- Jacobian : No
- Discontinuity Indicator : No (the transformation modules handles this part)
- Deadreckoning model : No
- Preferred Step Size : No (the step size is fixed at the beginning)
- Next Step Size : No (there is not an orchestrator for managing the step size and the step size are fixed)
- Order of Accuracy : No ( there is not extrapolation function)
- I/O Causality : Propagation Delay (the delay is used for the parallelization. However this delay is fixed during the simulation)
- Input Extrapolation : No (there is not extrapolation function)
- State Variables : Values
- Micro-Step Outputs : Yes (TVB and NEST give the output of each micro step but it can be modulated)
- Worst Case Execution Time : Yes (the worst case is when the minimum of delay is equal to the micro-time step (see Performance section))

##### Causality

Causal

##### Time Constraints

- Analytic Simulation : False (there does not analytic solution of this co-simulation)
- Scaled Real Time Simulation : Fixed for TVB and NEST
- Rollback Support : No (there is not rollback support for NEST and TVB)

##### Availability

local

#### Framework Requirements

- Standard: No standard (ad-hock solution)
- Coupling: Input/Output Assignments (A transformation modules between the two simulators take the role of the align of the I/O of the simulators)
- Number of Simulation Units : Two simulators
- Domain : Hybrid
- Dynamic structure : No (all the dependency is defined at the beginning)
- Co-simulation Rate : Single (unique size of step of synchronization between simulators and micro-step is fixed during the simulation)
- Communication Step Size : Fixed

- Strong Coupling Support : None – Explicit Method (the transformation module contains the information of the coupling of the simulators)
- Results Visualization : It can be in live or postmortem
- Communication Approach : Jacobi (however, the delays allow the separation of micro-steps without create errors)

##### Additional : characterization of the coupling[11][12]

The previous characterization is focusing more on the technical details but it is missing the characterization of the transformation modules. For the workflow of TVB-NEST, the scale are separate in space (micro- and macro-scale). The coupling between the simulators is a tightly coupling or cyclic coupling using a fix number of simulators instance. The workflows allows sequential or parallel execution depending of the number of initial conditions.

#### References

[1] Hahne, J. *et al.* NEST 3.0 (2021). URL https://zenodo.org/record/4739103.

[2] Sanz Leon, P. *et al.* The virtual brain: a simulator of primate brain network dynamics. *Front. Neuroinform.* **7** (2013). URL https://www.frontiersin.org/articles/10.3389/fninf.2013.00010/full. https://doi.org/10.3389/fninf.2013.00010 .

[3] Brette, R. & Gerstner, W. Adaptive exponential integrate-and-fire model as an effective description of neuronal activity. *Journal of Neurophysiology* **94** (5), 3637–3642 (2005). URL http://www.physiology.org/doi/10.1152/jn.00686.2005. https://doi.org/10.1152/jn.00686.2005 .

[4] Hagen, E. *et al.* Hybrid scheme for modeling local field potentials from point-neuron networks. *Cereb. Cortex* **26** (12), 4461–4496 (2016). URL https://academic.oup.com/cercor/article/26/12/4461/2333943. https://doi.org/10.1093/cercor/bhw237 .

[5] Shuman, T. *et al.* Breakdown of spatial coding and interneuron synchronization in epileptic mice. *Nat. Neurosci.* **23** (2), 229–238 (2020). URL https://www.ncbi.nlm.nih.gov/pmc/articles/PMC7259114/. https://doi.org/10.1038/s41593-019-0559-0 .

[6] Melozzi, F., Woodman, M. M., Jirsa, V. K. & Bernard, C. The virtual mouse brain: A computational neuroinformatics platform to study whole mouse brain dynamics **4** (3), ENEURO.0111–17.2017. URL http://www.eneuro.org/content/4/3/ENEURO.0111-17.2017. https://doi.org/10.1523/ENEURO.0111-17.2017 .

Springer Nature 2021 LATEX template



[7] Oh, S. W. *et al.* A mesoscale connectome of the mouse brain. *Nature* **508** (7495), 207–214 (2014). URL https://www.nature.com/articles/nature13186. https://doi.org/10.1038/nature13186 .

[8] di Volo, M., Romagnoni, A., Capone, C. & Destexhe, A. Biologically realistic mean-field models of conductance-based networks of spiking neurons with adaptation. *Neural Computation* **31** (4), 653–680 (2019). URL https://www.mitpressjournals.org/doi/10.1162/neco_a_01173. https://doi.org/10.1162/neco_a_01173 .

[9] Kuhn, A., Aertsen, A. & Rotter, S. Higher-order statistics of input ensembles and the response of simple model neurons. *Neural Computation* **15** (1), 67–101 (2003). URL http://www.mitpressjournals.org/doi/10.1162/089976603321043702. https://doi.org/10.1162/089976603321043702 .

[10] Gomes, C., Thule, C., Broman, D., Larsen, P. G. & Vangheluwe, H. Co-simulation: A survey. *Association for Computing Machinery* May 23, (2018). URL https://doi.org/10.1145/3179993.

[11] Chopard, B., Borgdorff, J. & Hoekstra, A. G. A framework for multi-scale modelling. *Philosophical Transactions of the Royal Society A: Mathematical, Physical and Engineering Sciences* **372** (2021), 20130378 (2014). URL https://royalsocietypublishing.org/doi/full/10.1098/rsta.2013.0378. https://doi.org/10.1098/rsta.2013.0378, publisher: Royal Society .

[12] Taveres-Cachat, E., Favoino, F., Loonen, R. & Goia, F. Ten questions concerning co-simulation for performance prediction of advanced building envelopes. *Building and Environment* **191**, 107570 (2021). URL https://www.sciencedirect.com/science/article/pii/S0360132320309379. https://doi.org/10.1016/j.buildenv.2020.107570 .
